# Restoration of Spermatogenesis is Dependent on Activation of a SPRY4-ERK Checkpoint Following Germline Stem Cell Damage

**DOI:** 10.1101/2025.10.12.681919

**Authors:** Ying Liu, Tansol Choi, Brad Pearson, Ryan Nachman, Whitney Woo, Na Xu, Ryan Schreiner, Romulo Hurtado, Marco Seandel, Shahin Rafii, Todd Evans

## Abstract

Mammalian spermatogonial stem cells (SSCs) sustain male fertility through continuous self-renewal and differentiation, leading to the production of haploid spermatozoa throughout adulthood. However, SSCs are vulnerable to genotoxic drugs, and patients receiving chemotherapy face a high risk of germline instability and infertility. The molecular mechanisms and cellular pathways that choreograph SSC recovery after chemotherapeutic insult remain unknown. Previously, we identified SPRY4 as an ERK-dependent negative feedback regulator of growth factor signaling that is critical for preservation of stem cell activity in cultured mouse SSCs. Here, we demonstrate that following alkylating agent busulfan (BU)-induced injury in adult mice, germline-specific *Spry4* gene deletion (*Spry4*^G-KO^) reduces stem cell regeneration with an enhanced genotoxic stress response and differentiation with rapidly enhanced nuclear ERK1/2 activity in undifferentiated (A_undiff_) spermatogonia (including SSCs). Genes essential for stem cell maintenance, including *Id1* and *Cxcl12*, were dysregulated by loss of *Spry4*. Furthermore, the MEK1/2 inhibitor PD0325901, but not mTORC1 inhibitor Rapamycin, was sufficient to promote spermatogonial proliferation in *Spry4*^G-KO^ testis 10 days post-BU treatment. Notably, the restoration of both spermatogonia pool and fertility was delayed in adult *Spry4*^G-KO^ males long-term after injury. In summary, germline-specific deletion of *Spry4* results in hyper-activation of the MAPK/ERK pathway in A_undiff_ spermatogonia, reducing spermatogonial genome integrity, unleashing excessive spermatogenesis after germline damage, and ultimately impairing germline regeneration in adult males. Our study indicates an essential role for SPRY4-ERK signaling as a molecular checkpoint in securing SSC recovery upon chemotherapy drug-induced germline damage, revealing how stem cells normally withstand environmental stress.

## Introduction

In adult mammalian testis, haploid spermatozoa are produced from unipotent spermatogonial stem cells (SSCs) through a tightly regulated process known as spermatogenesis. SSCs reside in the Sertoli cell-based niche at the basal lamina of the seminiferous tubules and are capable of both self-renewal and differentiation to maintain the germline stem cell pool while generating differentiated progeny throughout adulthood. In the mouse testis, SSC activity is enriched within a heterogenous population of undifferentiated spermatogonia (A_undiff_) composed of type A-single (A_s_) spermatogonia, A_paired_ (A_pr_) spermatogonia consisting of two interconnected cells, and A_aligned_ (A_al_) spermatogonia organized as chains of 4, 8, or 16 interconnected cells, with A_al_ spermatogonia ultimately transitioning into differentiated spermatogonia (A_diff_) and proceeding to meiosis [1, 2]. Accumulating evidence from lineage-tracing and live-imaging studies indicates that SSCs are not restricted to a fixed morphological state, but dynamically and reversibly transition between A_s_, A_pr_, and short A_al_ chains, under both homeostasis and regenerative conditions, highlighting the developmental plasticity within A_undiff_ spermatogonia [3–7].

Glial cell line-derived neurotrophic factor (GDNF) and fibroblast growth factor (FGF) are essential for SSC self-renewal and maintenance [8]. Both bind to receptor tyrosine kinases (RTKs) on the SSC membrane to activate the downstream effector Ras and initiate intracellular signaling cascades including the PI3K/AKT and MAPK/ERK pathways [9]. Both GDNF and FGF signaling regulate mTORC1, the central signaling hub that governs SSC fate commitment [8, 10]. In adult mouse testis, mTORC1 activity is suppressed in most self-renewing spermatogonia but induced by the PI3K/AKT pathway to generate cell-cycle-activated progenitors [11–13]. The PI3K/AKT/mTOR signaling pathway has been extensively explored as the master regulator that drives SSC proliferation and differentiation in both cultured SSCs and mouse models [14–16], although an in vitro study suggests that the MAPK/ERK pathway is also required to efficiently activate mTORC1 [17]. The MAPK/ERK pathway can be rapidly activated in response to various extracellular or intracellular stimuli to regulate distinct cellular processes including proliferation, differentiation, and regeneration [18]. Following sequential phosphorylation along the Ras/Raf/MEK/ERK signaling cascade, phosphorylated ERK1/2 (p-ERK1/2) impacts a broad set of substrates localized within the cytoplasm, including mTORC1 [19]. Activated p-ERK1/2 can also rapidly translocate from the cytoplasm to the nucleus to regulate transcriptional activity. The subcellular localization bestows p-ERK1/2 with distinct functions in regulating cellular responses to stimuli [20, 21]. Unlike PI3K/AKT and mTORC1, the function of the MAPK/ERK pathway in SSCs remains elusive. Several studies reported that both GDNF and FGF2 can activate the MAPK/ERK pathway and promote SSCs proliferation in vitro [15, 22, 23], although FGF2 has also been suggested to promote cultured SSCs to a more differentiation-primed state [24]. Studies of A_undiff_ spermatogonia within adult mouse testis indicate that intrinsic MAPK/ERK pathway components and regulators play a role in A_undiff_ spermatogonia self-renewal [25, 26], although MAPK/ERK signaling from the somatic niche has also been shown to contribute to the cyclical activity of SSCs [26].

Like many types of adult stem cells, SSCs are very sensitive to chemotherapy drugs including the alkylating agent busulfan (BU), which is particularly toxic to SSCs and other rapidly dividing spermatogonia [27, 28]. However, a few A_undiff_ spermatogonia can escape from low-dose BU treatment, repopulate rapidly after genotoxic stress, regenerate the seminiferous epithelium, and restore long-term fertility [29]. Although the physiological and molecular features of regenerative spermatogonia have been characterized in adult mice [30–32], how the germline-intrinsic signaling modulators respond to injury and cooperate to restore germline homeostasis remains obscure.

We identified SPRY4 (Sprouty4), primarily known as a negative feedback regulator of the RTK-MAPK/ERK pathway through its interference with Ras activation [33, 34], as a hallmark protein that is exclusively expressed in A_undiff_ spermatogonia within adult mouse germ cells [35]. In cultured SSCs, SPRY4 modulates the cellular response to FGF2 or GDNF through the MAPK/ERK pathway, with *Spry4* deletion enhancing early differentiation and diminishing stem cell activity in transplantation analysis. However, germline-specific *Spry4* deletion from adult mice (*Spry4*^G-KO^) does not affect stem cell colonization and differentiation when tested by transplanting total testicular cells to wild-type recipients [35]. To further address the SPRY4 function in the regeneration of SSCs in vivo, we employed a BU-induced germline damage and recovery mouse model to uncover the signaling pathways choreographing stem cell homeostasis after injury.

Toward this end, adult *Spry4*^G-KO^ or wild-type (*Spry4*^WT^) males were treated with a single low-dose of BU, and SSC recovery and differentiation were temporally and spatially determined. We found that quiescent SSCs within *Spry4*^WT^ mouse testis recovered from injury through a series of well-orchestrated transitions to achieve cell cycle activation and fate commitment. Notably, we found that germline-intrinsic SPRY4-ERK signaling must be carefully titrated to appropriately initiate stem cell regeneration. However, loss of *Spry4* resulted in hyperactivation of the MAPK/ERK pathway in A_undiff_ spermatogonia, enhanced genotoxic stress response and transient stem cell proliferation rapidly after damage, and unleashed spermatogonial differentiation. The transition from proliferating to fate-primed A_undiff_ spermatogonia was correlated with p-ERK1/2 translocation from the cytoplasm to the nucleus. We also show that young adult *Spry4*^G-KO^ males experience a fertility defect with delayed recovery of A_undiff_ spermatogonia pool after BU treatment, suggesting that dysregulation of the MAPK/ERK pathway is detrimental to SSC homeostasis during regeneration. Therefore, germline-intrinsic SPRY4-ERK signaling is essential to maintain and prevent exhaustion of SSCs after injury, thereby ensuring long-term spermatogenesis under environmental stresses through adulthood.

## Results

### Adult males with short-term germline-specific *Spry4* deletion can undergo spermatogenesis at steady state

We previously demonstrated that mouse SSCs respond robustly to GDNF or FGF2 induction in culture and activate *Spry4* expression to modulate the MAPK/ERK pathway [35]. Loss of *Spry4* promoted in vitro differentiation of SSCs at the expense of self-renewal. To investigate the in vivo cell autonomous function of SPRY4 under normal conditions, specifically in adult mice, we crossed *<u>G</u>fr*α*1-<u>C</u>reER^T2^;<u>R</u>osa26-LSL-td<u>T</u>omato* (referred to as GCRT) mice with *Spry4^flox/flox^* mice to create GCRT;*Spry4^flox/flox^*mice [35]. Germline-specific *Spry4* deletion (referred to as *Spry4*^G-KO^) was generated by injecting tamoxifen (100 mg/kg) intraperitoneally for two weeks in adult (2- to 4-month-old) GCRT;*Spry4^flox/flox^* mice. Tamoxifen effectively activates *Cre* in spermatogonia, which then pass the genetic modification to all stages of differentiating germline descendants in the testis [4]. As expected, robust expression of tdTomato, a stop-floxed lineage tracer, was observed within the *Spry4*^G-KO^ mouse testis (**Figure S1A**). Whole-mount immunofluorescence (IF) staining revealed that these tdTomato^+^ cells expressed many germ cell markers, including VASA (marking all germ cells), GFRα1(delineating the most primitive set of A_undiff_ spermatogonia), and MCAM/CD146 (delineating both A_undiff_ spermatogonia and progenitors [36, 37]), but not Vimentin (marking Sertoli cells, the somatic niche of spermatogonia). These data illustrate efficient gene deletion of *Spry4* within different stages of germ cells (**Figure S1B – 1D**).

We then collected testes for analysis from *Spry4*^G-KO^ or control *Spry4^flox/flox^* (referred to as *Spry4*^WT^) mice 6 weeks after the first tamoxifen injection (harvested within 4 months of age) for analysis. No marked difference in testis size (**Figure S2A**) or mature spermatozoa count (**Figure S2B**) was identified between *Spry4*^G-KO^ and *Spry4*^WT^ testes. Histological analysis revealed that seminiferous tubules in the testis sections from both *Spry4*^G-KO^ and *Spry4*^WT^ mice were populated with germ cells at different developmental stages, and functionally mature spermatozoa were identified in the cauda epididymis of both genotypes (**Figure S2C**). Taken together, GFRα1-CreER^T2^ induced germline-specific *Spry4* deletion does not significantly affect spermatogenesis in adult mouse testis in the short-term (6 weeks post-tamoxifen), suggesting that SPRY4 is dispensable for spermatogonia differentiation at steady state in young adulthood.

### *Spry4* deletion reduces stem cell genome integrity and regeneration in chemotherapy damaged adult mouse testis

To investigate Spry4 function in accelerated stem cell regeneration in vivo, we used a chemotherapy-induced germline damage and recovery model to explore whether loss of *Spry4* impacts stem cell activity during germline recovery [32]. The alkylating agent busulfan (BU) has been reported to cause apoptosis of undifferentiated (A_undiff_) and differentiating (A_diff_) spermatogonia in rodents [38]. Although the highest non-lethal BU dose (40 mg/kg) results in infertility with few spermatogonia remaining in the seminiferous tubules, lower doses of BU (e.g., 10 mg/kg) preserve some A_undiff_ spermatogonia that can restore the germline, providing a reliable model of spermatogenesis that requires SSC regeneration and differentiation [32, 38]. Therefore, we used a single low-dose of BU (10 mg/kg) to induce germline damage in adult *Spry4*^G-KO^ or *Spry4*^WT^ males and analyzed testes after recovering from BU treatment for 10 days (BU-D10), a latency period sufficient for surviving A_undiff_ spermatogonia to actively proliferate and initiate regeneration (**Figure 1A**).

**Figure 1.**
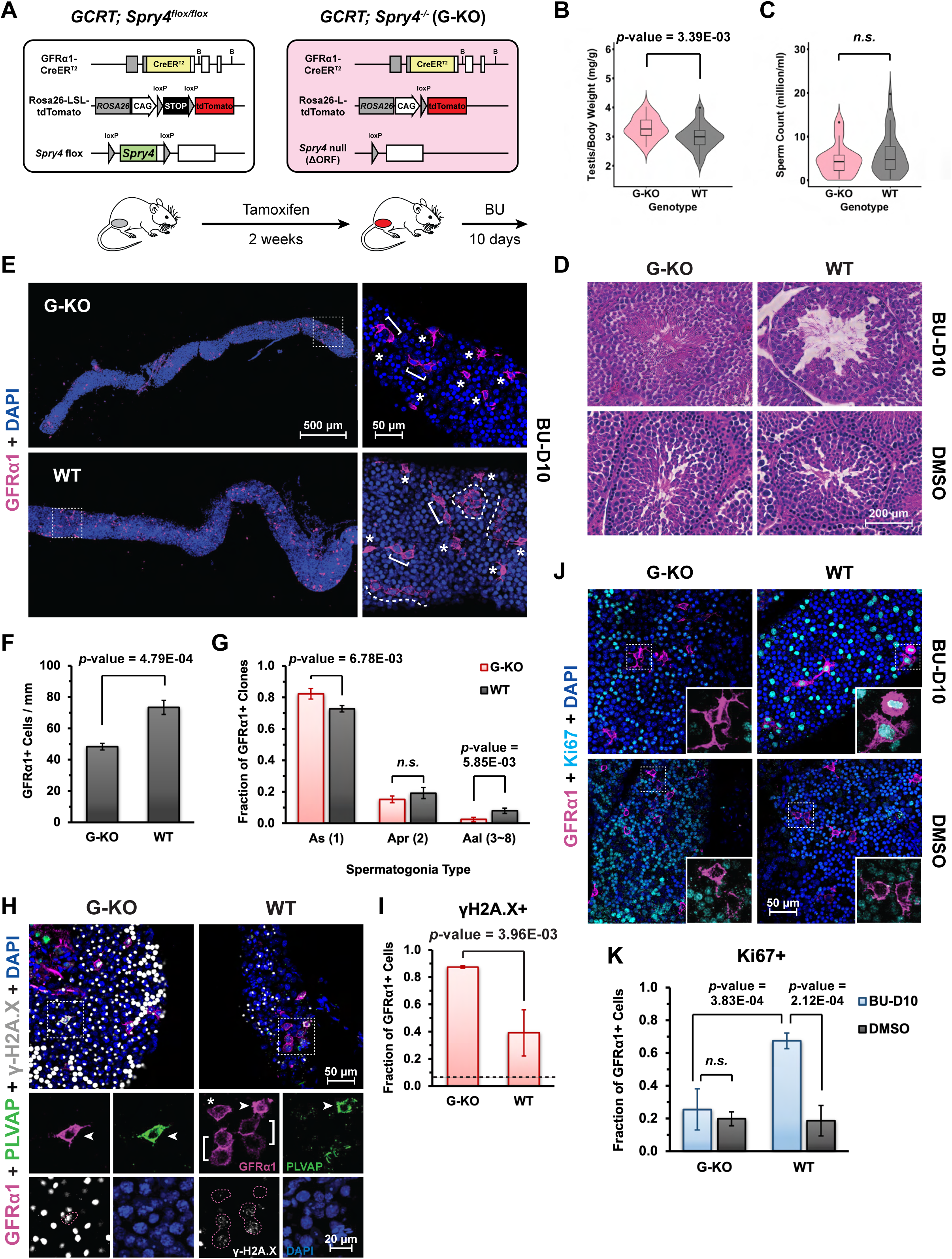
Following BU-induced damage, germline-specific *Spry4* deletion reduces A_undiff_ spermatogonia genome integrity and proliferation in adult mouse testis. **(A)** Schema illustrating the induction of germline-specific *Spry4* deletion and germline recovery in response to low-dose BU treatment in adult mouse testis. GCRT, *<u>Gfr</u>*<u>α*1*</u>*-<u>C</u>reER^T2^;<u>R</u>osa26-LSL-td<u>T</u>omato*. Testes collected from adult G-KO or WT mice 10 days post-BU treatment were compared for **(B)** testis to body weight ratio and **(C)** sperm count (n>10 mice analyzed per genotype). **(D)** Hematoxylin and eosin (H&E)–stained histological cross-sections of testes collected from adult *Spry4*^G-KO^ (G-KO) or *Spry4*^WT^ (WT) mice at 10 days post BU (BU-D10, Top) or post DMSO (DMSO, Bottom) treatment. Lower magnification images are shown in **Fig S3**. **(E)** Representative whole-mount immunofluorescence (IF) images illustrating GFRα1^+^ populations in tubules from *Spry4*^G-KO^ and *Spry4*^WT^ mice at BU-D10. Higher magnification images of indicated regions are shown on the right. Magenta, GFRα1; Blue, DAPI. Asterisks, A_s_ spermatogonia; Brackets, A_pr_ spermatogonia; Dashed lines, A_al_ spermatogonia (4 – 8 cells). **(F)** Quantification of GFRα1^+^ cells per mm of seminiferous tubules. **(G)** Fraction of A_s_, A_pr_, and A_al_ spermatogonia based on whole-mount IF analysis of GFRα1^+^ clones. Numbers in round brackets indicate cell numbers per GFRα1^+^ clone (n>3 mice analyzed per genotype). **(H)** Whole-mount IF staining on γH2A.X foci within the nuclei of A_undiff_ spermatogonia collected at BU-D10. Higher magnification images of the indicated regions are shown below. Magenta, GFRα1; Green, PLVAP; Grey, γH2A.X; Blue, DAPI. Arrowhead, GFRα1^+^PLVAP^+^ A_s_ spermatogonia; Asterisk, GFRα1^+^PLVAP^-^ A_s_ spermatogonia; Brackets, A_pr_ spermatogonia; Dashed outline, GFRα1^+^ A_s_ spermatogonia. **(I)** Quantification of γH2A.X^+^ cells within GFRα1^+^ cells in whole-mount IF. Dashed line, fraction of γH2A.X^+^ cells within GFRα1^+^ cells at steady state in wild-type mouse testis. (n>3 mice analyzed per condition, n>100 GFRα1^+^ cells scored per animal). **(J)** Whole-mount IF staining of the proliferation state in A_undiff_ spermatogonia collected after BU-D10 or DMSO treatment. Magenta, GFRα1; Cyan, Ki67; Blue, DAPI. Inset, higher magnification images of indicated regions without DAPI. **(K)** Quantification of Ki67^+^ cells within GFRα1^+^ cells in whole-mount IF staining of the tubules collected after BU-D10 (blue) or DMSO (black) treatment (n>4 mice analyzed per condition, n>100 GFRα1^+^ cells scored per animal). Data are mean (SD). *p*-value is calculated by unpaired one-tailed Student’s *t*-test. *n.s.*, not significant (*p*-value >= 0.05).

We found that testes collected from *Spry4*^G-KO^ males were significantly larger than those from *Spry4*^WT^ males at BU-D10 (**Figure 1B**), although both *Spry4*^G-KO^ and *Spry4*^WT^ mice generated similar sperm counts (**Figure 1C**). Given that the epididymal sperm collected at BU-D10 most likely originated from post-meiotic germ cells at the time of BU exposure, these comparable sperm counts suggest that SPRY4 does not affect the survival or maturation of post-meiotic germ cells following chemotherapy treatment. Histological analysis of the testis sections collected at BU-D10 showed that the seminiferous tubules within *Spry4*^G-KO^ mouse testes contained germ cells at different developmental stages, similar to the tubules of *Spry4*^G-KO^ or *Spry4*^WT^ mice with control DMSO treatment (DMSO) (**Figure 1D** and **S3**). In contrast, most seminiferous tubules in the testis sections of *Spry4*^WT^ mice were degenerated with considerable loss of developing germ cells and mature spermatozoa, a phenotype which has been reported in low-dose BU treated males [38] (**Figure 1D** and **S3**).

By whole-mount IF staining of seminiferous tubules, we identified GFRα1^+^ cells in testes collected from both *Spry4*^G-KO^ and *Spry4*^WT^ mice at BU-D10 (**Figure 1E**). Comparing to *Spry4*^WT^ mice post-BU, the tubules collected from *Spry4*^G-KO^ mice contained less GFRα1^+^ cells (**Figure 1F**), with over 82% of GFRα1^+^ cells existing as A_s_ spermatogonia, the most primitive spermatogonia resembling SSCs (**Figure 1E**, asterisks). The fractions of A_s_ spermatogonia within the *Spry4*^G-KO^ tubules were significantly higher than those identified in the tubules collected from *Spry4*^WT^ mice (**Figure 1G**). A_s_ spermatogonia can divide to form two spermatogonia connected by an intercellular bridge (A_pr_), then proceed to form a chain of aligned (A_al_) spermatogonia. We identified over 15% of A_pr_ spermatogonia clones and numerous chains of A_al_ spermatogonia containing up to 8 GFRα1^+^ cells within *Spry4*^WT^ tubules (**Figure 1E** and **1G**, WT). However, A_al_ spermatogonia chains with 8 GFRα1^+^ cells were barely detected within the tubules collected from *Spry4*^G-KO^ mice (**Figure 1E** and **1G**, G-KO).

A cell-surface protein PLVAP was reported as a marker of the most primitive SSCs that co-express GFRα1 mainly in A_s_ spermatogonia [7]. We found that the fractions of PLVAP^+^ cells within single GFRα1^+^ cells remain similar between *Spry4*^G-KO^ and *Spry4*^WT^ mice at BU-D10, suggesting that deleting SPRY4 does not affect the transition between renewal-biased and differentiation-primed states within A_s_ spermatogonia at the beginning of regeneration (Figure **S4**). Notably, γH2A.X foci, a key biomarker for accumulated DNA double-strand breaks and cellular senescence, were identified in over **80%** of GFRα1^+^ cells (including both PLVAP^+^ and PLVAP^-^ cells) within the *Spry4*^G-KO^ tubules at BU-D10, remarkably higher than those identified in the *Spry4*^WT^ mice (∼**40%** of GFRα1^+^ cells) (**Figure 1H** and **1I**).

To evaluate regenerative capacity in A_undiff_ spermatogonia, we co-stained cell proliferation marker Ki67 with GFRα1. We found that only about 20% of GFRα1^+^ cells were positive for Ki67 in the tubules collected from *Spry4*^G-KO^ or *Spry4*^WT^ mice without chemotherapeutic insult (DMSO), suggesting that most A_undiff_ spermatogonia remain quiescent without damage (**Figure 1J** and **1K**, DMSO). However, nearly 70% of *Spry4*^WT^ GFRα1^+^ cells surviving BU treatment were highly proliferative with strong Ki67 staining at BU-D10 (**Figure 1J** and **1K**, WT, BU-D10), as has been reported in regenerating spermatogonia [32]. Notably, only about 25% of *Spry4*^G-KO^ GFRα1^+^ cells were actively proliferating at BU-D10, suggesting that spermatogonia regeneration is repressed in *Spry4*^G-KO^ mice after chemotherapy drug-induced germline damage (**Figure 1J** and **1K**, G-KO, BU-D10).

### *Spry4*^G-KO^ spermatogonia exhibit enhanced differentiation during germline recovery

To characterize spermatogonia populations within the seminiferous tubules of BU-damaged testis, we applied whole-mount IF staining for MCAM to detect both A_undiff_ and A_diff_ spermatogonia within the tubules collected from *Spry4*^G-KO^ or *Spry4*^WT^ mouse testes at BU-D10. Although more GFRα1^+^ cells (A_undiff_ spermatogonia) were found in the tubules of *Spry4*^WT^ mice than in *Spry4*^G-KO^ mice (**Figure 1F**), MCAM^+^ cells were sparse in multiple tubules collected from *Spry4*^WT^ mice, with some regions completely devoid of GFRα1^-^MCAM^+^ cells (A_diff_ spermatogonia), corresponding with the initiation of regeneration post damage (**Figure 2A**, WT). Strikingly, the tubules collected from *Spry4*^G-KO^ mice were nearly entirely populated with GFRα1^-^MCAM^+^ cells (**Figure 2A**, G-KO). Similar to the GFRα1^+^ A_undiff_ spermatogonia, most MCAM^+^ cells were quiescent in *Spry4*^G-KO^ testes (**Figure S5**). Further investigation with RARγ (marking differentiation-destined progenitors) and c-KIT (marking A_diff_ spermatogonia) by flow cytometry analysis 4 days after busulfan treatment (BU-D4) revealed a similar reduction of c-KIT^+^ cells in both genotypes, indicating that A_diff_ spermatogonia do not acquire SPRY4-dependent sensitivity to busulfan early post-injury (**Figure S6A – C**). However, the RARγ^+^c-KIT^-^ population is markedly enriched in *Spry4*^G-KO^ testes compared with *Spry4*^WT^ controls at BU-D4, with RARγ^+^ progenitor cells highly proliferative in *Spry4*^G-KO^ testes but quiescent in *Spry4*^WT^ testes (**Figure S6D – G**). Thus, following comparable early depletion of A_diff_ spermatogonia (c-KIT^+^), SPRY4-deficient progenitors (RARγ^+^) prematurely re-enter the cell cycle at BU-D4, leading to precocious spermatogonial differentiation at BU-D10.

**Figure 2.**
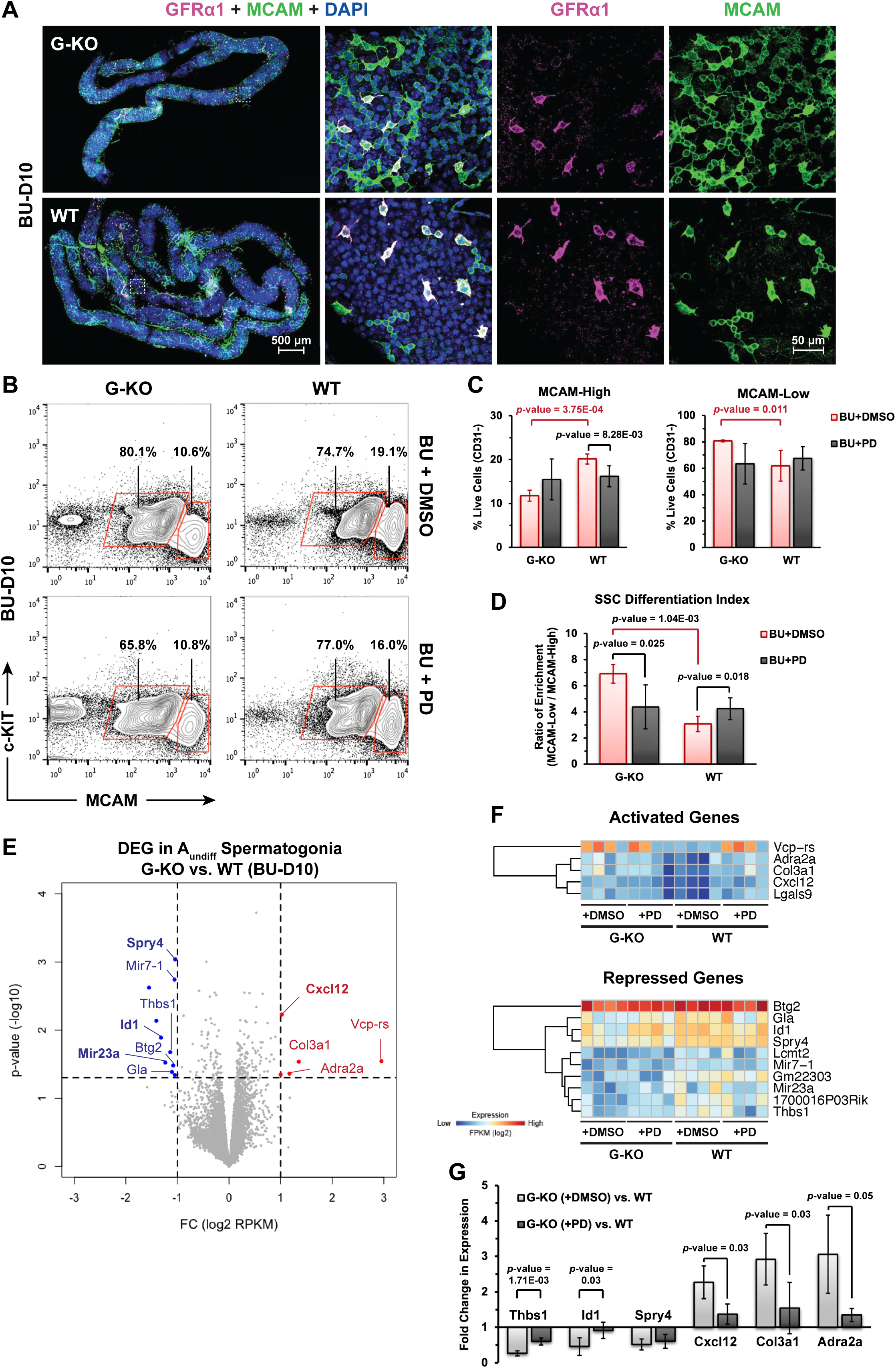
*Spry4*^G-KO^ spermatogonia exhibit enhanced differentiation with transcriptional changes at selected genes during germline regeneration (BU-D10). **(A)** Representative whole-mount IF images illustrating GFRα1^+^ and MCAM^+^ populations in tubules from *Spry4*^G-KO^ and *Spry4*^WT^ mice at BU-D10. Higher magnification images of indicated regions are shown on the right. Magenta, GFRα1; Green, MCAM; Blue, DAPI. **(B)** Fluorescence-activated cell sorting (FACS) strategy for isolation of MCAM^High^c-KIT^-^ (A_undiff_) and MCAM^Low^ (A_diff_) spermatogonia. **(C)** Percentage of cells within gates are indicated (n>3 mice analyzed per condition). **(D)** <u>S</u>SC <u>D</u>ifferentiation Index (SDI) calculated by the ratio of enrichment between MCAM^Low^ (A_diff_) and MCAM^High^c-KIT^-^ (A_undiff_) spermatogonia seen in Fig. 2C. **(E)** Volcano plot of expression profiles comparing MCAM^High^c-KIT^-^ population isolated from *Spry4*^G-KO^ or *Spry4*^WT^ mouse testes at BU-D10. Selected genes were highlighted based on expression changes: Red, genes upregulated in *Spry4*^G-KO^ mice; Blue, genes downregulated in *Spry4*^G-KO^ mice. (n=4 mice analyzed per condition) **(F)** Expression of genes activated or suppressed in *Spry4*^G-KO^ mice. Red and blue indicate relatively high and low gene expression, respectively. **(G)** qRT-PCR analysis of selected genes as highlighted in Fig. 2E (n=4 mice analyzed per condition). Data are mean (SD). *p*-value is calculated by unpaired one-tailed Student’s *t*-test.

We further quantified the spermatogonia by fluorescence-activated cell sorting (FACS) of MCAM^+^ cells, with the high MCAM and negative c-KIT (MCAM^High^c-KIT^-^) population enriched for A_undiff_ spermatogonia and low MCAM (MCAM^Low^) population enriched for A_diff_ spermatogonia (**Figure S7A – C**). We observed tdTomato signal in most MCAM^High^c-KIT^-^ cells isolated from *Spry4*^G-KO^ mouse testes, confirming efficient GFRα1-Cre^ERT2^-induced recombination within A_undiff_ spermatogonia (**Figure S7D**). Notably, the MCAM^High^c-KIT^-^population isolated from *Spry4*^G-KO^ mice was significantly reduced compared to those isolated from *Spry4*^WT^ mice at BU-D10 (**Figure 2B** and **2C**, MCAM-High, BU+DMSO). Compared with *Spry4*^WT^ mice, the MCAM^Low^ population containing more differentiating spermatogonia was significantly increased in *Spry4*^G-KO^ mouse testes (**Figure 2B** and **2C**, MCAM-Low, BU+DMSO). To quantify differentiation status in testicular cells, we defined the <u>S</u>SC <u>D</u>ifferentiation Index (SDI) by calculating the ratio of enrichment between MCAM^Low^ (A_diff_) and MCAM^High^c-KIT^-^ (A_undiff_) spermatogonia (See **Methods**). The SDI indicated that the *Spry4*^G-KO^ spermatogonia retain a more differentiated status compared to the *Spry4*^WT^ spermatogonia at BU-D10 (**Figure 2D**, BU+DMSO).

SPRY4 has been identified as a negative feedback regulator of the MAPK/ERK pathway. Our prior studies with cultured adult mouse SSCs revealed that SPRY4 is critical for preserving stem cell activity through ERK-dependent regulation of FGF and GDNF signaling. To investigate the function of SPRY4-ERK signaling during chemotherapy-induced germline damage and recovery in vivo, ERK activity in *Spry4*^G-KO^ or *Spry4*^WT^ mice was blocked by treating mice daily with the MEK inhibitor PD0325901 (PD, also known as mirdametinib) prior to regeneration (3 days after low-dose BU treatment) and analyzed at BU-D10. Notably, PD treatment resulted in similar MCAM^High^c-KIT^-^ or MCAM^Low^ populations in *Spry4*^G-KO^ and *Spry4*^WT^ mice, while the MCAM^High^c-KIT^-^ population collected from *Spry4*^WT^ mice was slightly reduced with PD treatment (**Figure 2B** and **2C**, BU+PD). Corresponding with these findings, the SDI of PD-treated *Spry4*^G-KO^ spermatogonia is substantially reduced to the levels seen in *Spry4*^WT^ spermatogonia, suggesting that PD treatment can restore the *Spry4*^G-KO^ spermatogonia to a less differentiated status (**Figure 2D**, BU+PD). Quantification of the GFRα1^+^ cells (A_undiff_ spermatogonia) within the whole-mount IF stained seminiferous tubules confirmed that inhibiting the MAPK/ERK pathway by PD treatment does not affect the maintenance and clone composition of regenerating A_undiff_ spermatogonia in *Spry4*^G-KO^ mice (**Figure S8**). Thus, rather than killing A_undiff_ spermatogonia in *Spry4*^G-KO^ mice, PD treatment appears to promote their developmental dynamics toward A_diff_ spermatogonia.

### Characterization of SPRY4-dependent genes in regenerating spermatogonia

To investigate the molecular mechanism underlying spermatogonia hyper-differentiation in *Spry4*^G-KO^ mice post-BU, we applied bulk RNA-seq to characterize *Spry4*-dependent genes in MCAM^High^c-KIT^-^ (A_undiff_) spermatogonia at BU-D10. Surprisingly, very few genes demonstrated significant differential expression patterns (p-value < 0.05 and |fold change of log2 FPKM| > 1) in the A_undiff_ spermatogonia FACS-isolated from *Spry4*^G-KO^ mouse testes compared to *Spry4*^WT^ (**Figure 2E**). As expected, transcripts for the *Spry4* gene were significantly reduced in the A_undiff_ spermatogonia collected from *Spry4*^G-KO^ mouse testes (**Figure 2E – 2G**). Although PD treatment down-regulated *Spry4* gene expression in the A_undiff_ spermatogonia collected from *Spry4*^WT^ mouse testes (**Figure S5**), reflecting the role of SPRY4 as a MAPK/ERK-dependent negative feedback regulator, no expression difference was identified from *Spry4*^G-KO^ spermatogonia with or without PD treatment (**Figure 2F, 2G,** and **S5**). Among the genes downregulated after *Spry4* deletion, *Id1* is critical for mammalian stem cell maintenance by promoting quiescence and preventing premature differentiation in multiple adult stem cells [39]. Notably, treating *Spry4*^G-KO^ mice with the MAPK/ERK pathway inhibitor PD restored *Id1* gene expression levels to those seen in *Spry4*^WT^ mice (**Figure 2F, 2G,** and **S5**). Besides the *Id1* gene, two microRNA genes, Mir7-1 and Mir23a, were downregulated in *Spry4*^G-KO^ spermatogonia (**Figure 2E, 2F**). Mir7-1 is one of three genomic loci encoding miR-7, a highly conserved miRNA playing key roles in both normal organ development and tumorigenesis [40]. Several studies in male subfertile patients indicate that the levels of mature guide strand of Mir23a may associate with spermatogenesis-related gene expression [41, 42]. However, the functions of both microRNAs have not been investigated in SSCs. We also found that the expression of *Cxcl12*, a chemokine previously reported as a niche-derived factor associated with spermatogonial migration [43, 44], was significantly upregulated in *Spry4*^G-KO^ A_undiff_ spermatogonia, with this effect largely diminished after PD treatment (**Figure 2E – 2G,** and **S9A**). To further explore SPRY4–ERK-regulated transcriptional programs in vivo, we reanalyzed previously published single-cell RNA-seq (scRNA-seq) data from FACS-isolated *Spry4*^+^ (Venus^+^) A_undiff_ spermatogonia obtained from *Spry4*^H2B-Venus^ reporter mice [35]. This analysis revealed expression of several SPRY4-dependent genes identified in regenerating spermatogonia, including *Id1* and *Cxcl12*. Notably, the expressions of both genes were restricted in specific subsets of spermatogonial progenitor (*Id1* in Spg3, *Cxcl12* in Spg2) rather than broadly expressed across A_undiff_ spermatogonia, indicating a developmental stage-specific transcriptional response downstream of SPRY4–ERK signaling (**Figure S9B** and **C**).

### SPRY4 is required for A_undiff_ spermatogonia regeneration through inhibition of nuclear ERK activity

ERK1/2 can be activated by growth factors to p-ERK1/2 and translocate from cytoplasm to nucleus to modulate gene expression [21]. Our study with cultured SSCs revealed that SPRY4 is a critical regulator of MAPK/ERK signaling patterns during stem cell differentiation [35]. By whole-mount IF staining of *Spry4*^WT^ testicular tubules, we found that p-ERK1/2 was excluded from most spermatogonia, including GFRα1^+^ (A_undiff_) and GFRα1^-^MCAM^+^ (A_diff_) spermatogonia during regeneration (BU-D10) (**Figure 3A** and **S10A**, WT, BU+DMSO), with only about 10% of GFRα1^+^ spermatogonia slightly enriched for p-ERK1/2 in the cytoplasm or within the whole cells (**Figure 3B**, WT). By contrast, p-ERK1/2 was enriched in nearly 60% of GFRα1^+^ spermatogonia identified from *Spry4*^G-KO^ testicular tubules, and 47% of GFRα1^+^ spermatogonia contained p-ERK1/2 activity within the entire cell including the nuclei (**Figure 3A, 3B** and **S10B**, G-KO, BU+DMSO). Importantly, inhibiting the MAPK/ERK pathway with PD treatment was sufficient to eliminate p-ERK1/2 from the spermatogonia in both *Spry4*^G-KO^ and *Spry4*^WT^ testes, although p-ERK1/2 was still detected in many tdTomato^-^ or MCAM^-^ cells (Sertoli cells) within the seminiferous tubules (**Figure 3A** and **S10**, BU+PD, asterisks). Several studies have suggested that the MAPK/ERK pathway may regulate spermatogonia homeostasis through mTORC1, which has been reported as a critical inducer of A_undiff_ spermatogonia regeneration. Furthermore, inhibition of mTORC1 has been reported to trigger compensatory activation of the MAPK/ERK pathway via RPS6-PI3K-Ras-dependent feedback mechanisms in normal cell lines and human cancer models [45]. However, we did not observe significant p-ERK1/2 in the spermatogonia of either *Spry4*^G-KO^ or *Spry4*^WT^ mice treated with Rapamycin (RAPA), the inhibitor of mTORC1 (**Figure 3A** and **S10**, BU+RAPA).

**Figure 3.**
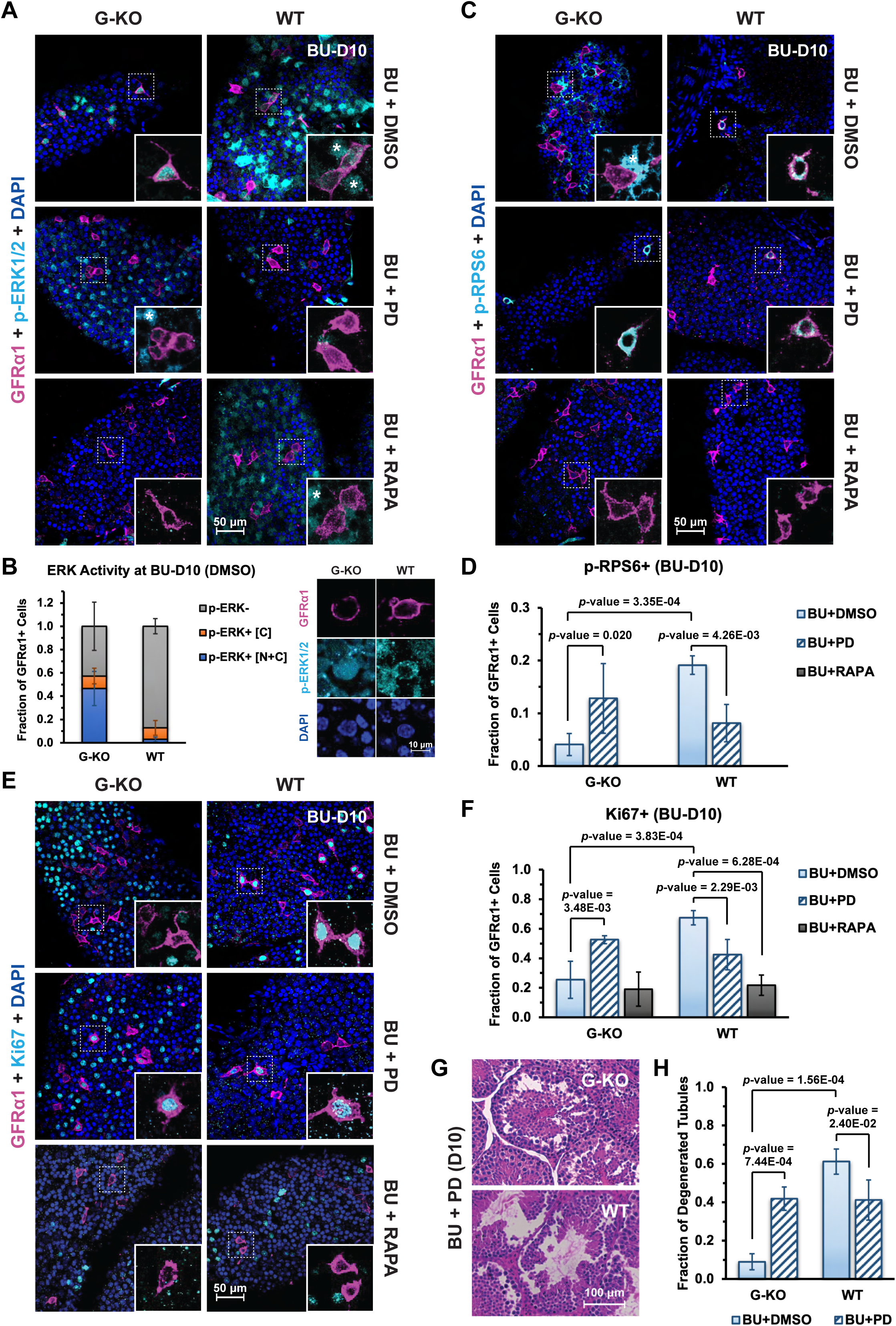
SPRY4 is required for A_undiff_ spermatogonia regeneration through inhibiting ERK signaling activity. Adult *Spry4*^G-KO^ or *Spry4*^WT^ mice were treated with a single low-dose BU and then daily with DMSO (BU+DMSO), PD0325901 (BU+PD), or Rapamycin (BU+RAPA) starting from day 3 post-BU. Testes were collected for analysis at 10 days post-BU by whole-mount IF staining for **(A)** ERK activity. Magenta, GFRα1; Cyan, p-ERK1/2; Blue, DAPI, or **(C)** mTORC1 activity. Magenta, GFRα1; Cyan, p-RPS6; Blue, DAPI, or **(E)** Proliferation. Magenta, GFRα1; Cyan, Ki67; Blue, DAPI. Inset, higher magnification images of indicated regions without DAPI. Asterisks, somatic cells. GFRα1^+^ cells in whole-mount IF were counted for the fraction of **(B)** p-ERK1/2 enrichment within both nucleus and cytoplasm (p-ERK^+^ [N+C]), only in the cytoplasm (p-ERK^+^ [C]), or no detectable p-ERK1/2 (p-ERK^-^) at BU-D10 (BU+DMSO), with higher magnification images of GFRα1^+^ cells in *Spry4*^G-KO^ or *Spry4*^WT^ tubules shown on the left, **(D)** p-RPS6^+^ cells, or **(F)** Ki67^+^ cells. n>3 mice analyzed per condition, n>100 GFRα1^+^ cells scored per animal. **(G)** H&E–stained histological cross-sections of adult mouse testes collected at 10 days post BU with PD0325901 treatment (BU+PD, D10). **(H)** Fraction of degenerated tubules identified in Fig. 3G (n=3 mice analyzed per condition). Data are mean (SD). *p*-value is calculated by unpaired one-tailed Student’s *t*-test.

To detect mTORC1 activity within spermatogonia during regeneration, we analyzed phosphorylated ribosomal protein S6 (p-RPS6) by whole-mount IF staining of the seminiferous tubules collected at BU-D10. Consistent with previous studies [32], we identified nearly 20% of GFRα1^+^ cells with strong cytoplasmic p-RPS6 in the tubules collected from *Spry4*^WT^ mice at BU-D10 (**Figure 3C** and **3D**, WT, BU+DMSO). Unexpectedly, mTORC1 activity was significantly reduced in *Spry4*^G-KO^ tubules, with only about 4% of GFRα1^+^ cells positive for p-RPS6 (4.7-fold less) at BU-D10 (**Figure 3C** and **3D**, G-KO, BU+DMSO). Furthermore, inhibition of ERK activity with PD treatment effectively increased the numbers of p-RPS6^+^ cells in *Spry4*^G-KO^ tubules to over 10% of GFRα1^+^ cells, comparable to that of *Spry4*^WT^ tubules after PD treatment (**Figure 3C** and **3D**, BU+PD). No p-RPS6^+^GFRα1^+^ cells were detected in *Spry4*^G-KO^ or *Spry4*^WT^ mice receiving both low-dose BU and mTORC1 inhibitor RAPA (**Figure 3C** and **3D**, BU+RAPA).

Activation of mTORC1 has been reported to be essential for the regenerative response in spermatogonia [32]. Correspondingly, quantification of Ki67^+^ cells by whole-mount IF staining of *Spry4*^WT^ tubules revealed that the number of proliferating A_undiff_ spermatogonia was reduced from nearly 70% of GFRα1^+^ cells in mice undergoing normal regeneration (BU+DMSO) to about 20% of GFRα1^+^ cells in mice receiving RAPA treatment (BU+RAPA) (**Figure 3E** and **3F**, WT). However, RAPA treatment did not affect proliferation of GFRα1^+^ cells in the tubules collected from *Spry4*^G-KO^ mice, further indicating that *Spry4* deletion inactivated mTORC1 in A_undiff_ spermatogonia at BU-D10 (**Figure 3E** and **3F**, G-KO, BU+RAPA). Notably, inhibition of MAPK/ERK pathway activity with PD treatment, while partially suppressing spermatogonial proliferation in *Spry4*^WT^ tubules, significantly restored spermatogonia regeneration in *Spry4*^G-KO^ tubules, shown by an increased fraction of Ki67^+^GFRα1^+^ cells from 25% (BU+DMSO) to over 50% (BU+PD) (**Figure 3E** and **3F**, G-KO). We also found that although BU treatment caused minimal damage in *Spry4*^G-KO^ tubules at BU-D10 (**Figure 1D**), PD treatment significantly impacted spermatogenesis with nearly 40% of tubules degenerated in both *Spry4*^G-KO^ and *Spry4*^WT^ mouse testes (**Figure 3G** and **3H**, BU+PD), suggesting that an active MAPK/ERK pathway is required for spermatogonia differentiation.

### Hyper-activated MAPK/ERK pathway enhances spermatogonia proliferation in *Spry4*^G-KO^ **testis shortly after damage**

Our *Spry4*^G-KO^ mouse model at BU-D10 revealed the importance of appropriate ERK activity in regulating spermatogonia recovery during regeneration. To assess effects of germline-specific *Spry4* deletion during the initial phase of regeneration, we analyzed the proliferation of A_undiff_ spermatogonia by whole-mount IF staining of seminiferous tubules collected at multiple time points. Remarkably, shortly after damage (2 days post-BU, BU-D2) the *Spry4*^G-KO^ tubules contained more Ki67^+^GFRα1^+^ cells than *Spry4*^WT^ tubules (**Figure 4A** and **4B**). Nearly 40% of GFRα1^+^ cells in the *Spry4*^G-KO^ tubules displayed strong Ki67 staining (Ki67^+^) until 6 days post-BU (BU-D6). By BU-D10, Ki67 decreased to a basal level (∼20%) of GFRα1^+^ cells, where it was maintained long-term after recovering from damage (up to 123 days post-BU) (**Figure 4A**, G-KO). Conversely, most GFRα1^+^ cells remained quiescent in *Spry4*^WT^ tubules at BU-D2, then gradually turned active from 4 days post-BU (BU-D4) until reaching the peak for proliferation at BU-D10 (**Figure 4A**, WT). Inhibition of ERK or mTORC1 activity with PD or RAPA treatment did not affect GFRα1^+^ cell proliferation in *Spry4*^WT^ tubules at BU-D2 (**Figure 4B** and **4C**, WT). However, both inhibitors reduced proliferation of the GFRα1^+^ cells in *Spry4*^G-KO^ tubules to similar levels that were comparable to those in *Spry4*^WT^ tubules (**Figure 4B** and **4C**, G-KO). In addition, robust cytoplasmic p-RPS6 was identified in the GFRα1^+^ cells in *Spry4*^G-KO^ tubules; this was effectively blocked by PD or RAPA treatment (**Figure 4D** and **4E**, G-KO). However, GFRα1^+^ cells in *Spry4*^WT^ tubules did not respond to either PD or RAPA treatment for enrichment of cytoplasmic p-RPS6 (**Figure 4D** and **4E**, WT).

**Figure 4.**
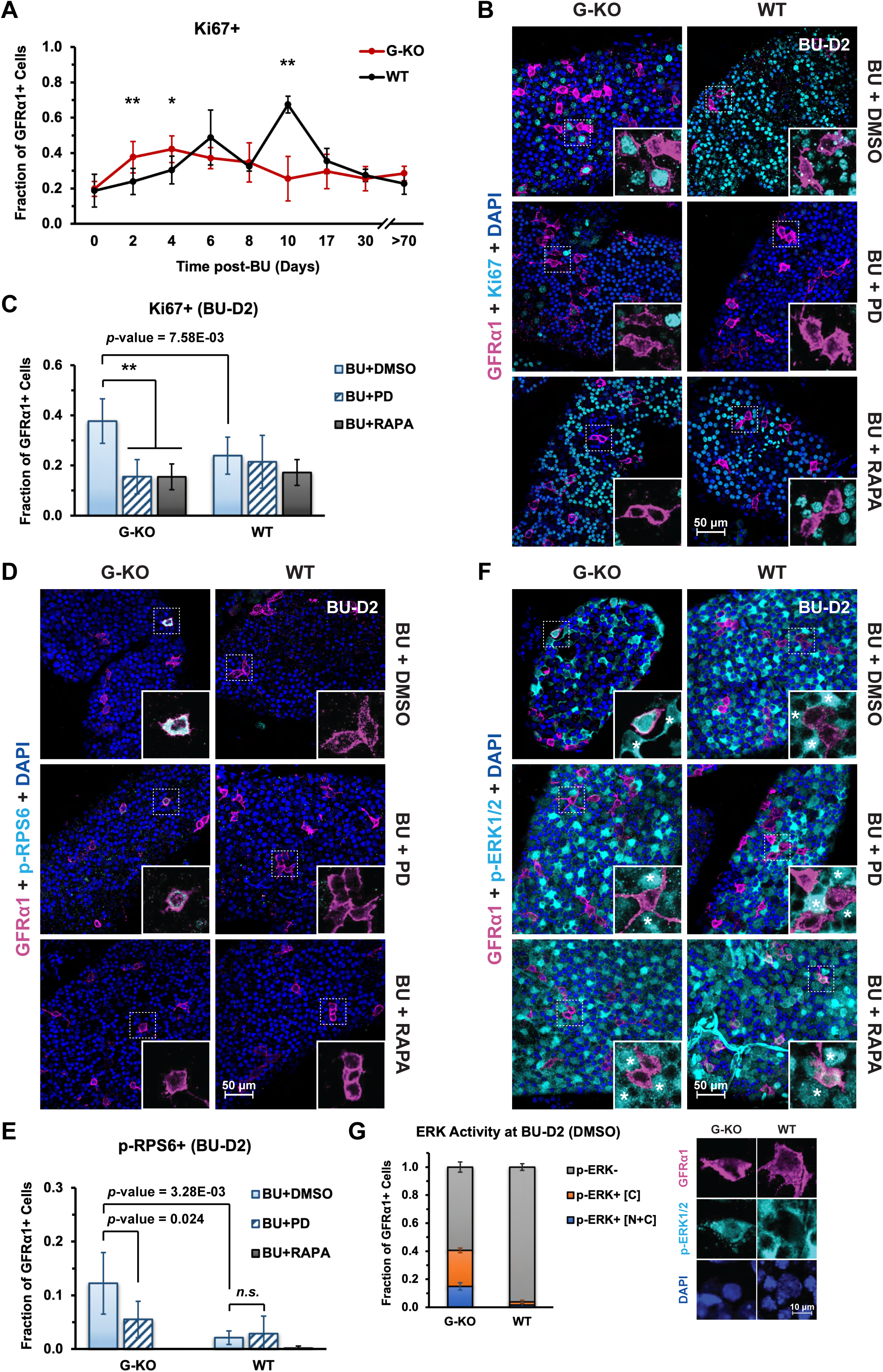
Hyper-activated ERK signaling enhances spermatogonia proliferation in *Spry4*^G-KO^ testis rapidly after damage (BU-D2). Adult *Spry4*^G-KO^ or *Spry4*^WT^ mice were treated with a single low-dose BU, and testes were collected at various time points post-BU as indicated. **(A)** Fraction of Ki67^+^ cells within GFRα1^+^ cells based on whole-mount IF. (n>3 mice analyzed per condition per time point, n>100 GFRα1^+^ cells scored per animal). Mice were treated with single low-dose BU together with DMSO (BU+DMSO), PD0325901 (BU+PD), or Rapamycin (BU+RAPA). Testes were collected 2 days post-BU (BU-D2) and analyzed by whole-mount IF staining for **(B)** Proliferation. Magenta, GFRα1; Cyan, Ki67; Blue, DAPI, **(D)** mTORC1 activity. Magenta, GFRα1; Cyan, p-RPS6; Blue, DAPI, or **(F)** ERK activity. Magenta, GFRα1; Cyan, p-ERK1/2; Blue, DAPI. Inset, higher magnification images of indicated regions without DAPI. Asterisks, somatic cells. GFRα1^+^ cells in whole-mount IF were counted for the fraction of **(C)** Ki67^+^ cells and **(E)** p-RPS6^+^ cells in the testes collected after indicated treatment, or **(G)** p-ERK1/2 enrichment within both nucleus and cytoplasm (p-ERK^+^ [N+C]), only in the cytoplasm (p-ERK^+^ [C]), or no detectable p-ERK1/2 (p-ERK^-^) at BU-D2 (BU+DMSO), with higher magnification images of GFRα1^+^ cells in *Spry4*^G-KO^ or *Spry4*^WT^ tubules shown on the left. n>3 mice analyzed per condition, n>100 GFRα1^+^ cells scored per animal. Data are mean (SD). *p*-valu*e* is calculated by unpaired one-tailed Student’s *t*-test. **, *p*-value < 0.01.

Notably, most tubules collected from *Spry4*^G-KO^ or *Spry4*^WT^ mice demonstrated profound p-ERK1/2 signaling in non-spermatogonial cells (MCAM^-^ or tdTomato^-^), revealing enhanced ERK activity within the somatic niche of germ cells shortly after damage (**Figure 4F** and **S11**, asterisks). p-ERK1/2 was identified within nearly 40% of GFRα1^+^ cells (26% mainly in cytoplasm and 14% within the entire cell) in *Spry4*^G-KO^ tubules but not in *Spry4*^WT^ tubules at BU-D2 (**Figure 4F** and **4G**, BU+DMSO). Both PD and RAPA reduced the enrichment of p-ERK1/2 within GFRα1^+^ cells in *Spry4*^G-KO^ tubules, although p-ERK1/2 was detected within a small number of GFRα1^+^ cells in *Spry4*^WT^ tubules after RAPA treatment (**Figure 4F**, BU+PD and BU+RAPA). These data suggest that although low-dose BU treatment does not induce a rapid regeneration response in *Spry4*^WT^ tubules, deletion of *Spry4* from spermatogonia results in hyperactive ERK right after damage. As such, stem cell proliferation is promoted rapidly yet transiently through ERK-mTORC1 signaling.

### SPRY4 is required for restoration of A_undiff_ spermatogonia and male fertility after damage

To explore the long-term consequences of germline-specific *Spry4* deletion on spermatogonia regeneration, we applied whole-mount IF to analyze GFRα1^+^ cells at multiple time points after BU treatment. Consistent with previous reports [32, 38, 46], the density of GFRα1^+^ cells in *Spry4*^WT^ tubules decreased at 4 days post-BU, then increased and transiently overshot baseline levels between 10 and 30 days post-BU, before recovering to steady-state levels after more than two months (72 ∼ 94 days post-BU) (**Figure 5A**). However, similar to the *Spry4*^G-KO^ tubules collected at BU-D10 (**Figure 1E** and **1F**), GFRα1^+^ cells remained sparsely located within the *Spry4*^G-KO^ tubules at 30 days post-BU (BU-D30), with the corresponding GFRα1^+^ cell densities lower than those collected from *Spry4*^WT^ tubules (**Figure 5A, S12A** and **S12B**). Furthermore, most GFRα1^+^ cells (∼80%) in the *Spry4*^G-KO^ tubules persisted as the most primitive A_s_ spermatogonia with intensely stained γH2A.X foci in nuclei, at a significantly higher rate than those identified in the *Spry4*^WT^ tubules at BU-D30 (**Figure S12C – E**). Conversely, the fractions of A_al_ spermatogonia were reduced in *Spry4*^G-KO^ tubules (**Figure S12C**). Notably, the number of GFRα1^+^ cells doubled within two months in *Spry4*^G-KO^ tubules after BU-D30 and overshot between 72 and 94 days post-BU, suggesting a delayed spermatogonia regeneration with *Spry4* deletion (**Figure 5A**).

**Figure 5.**
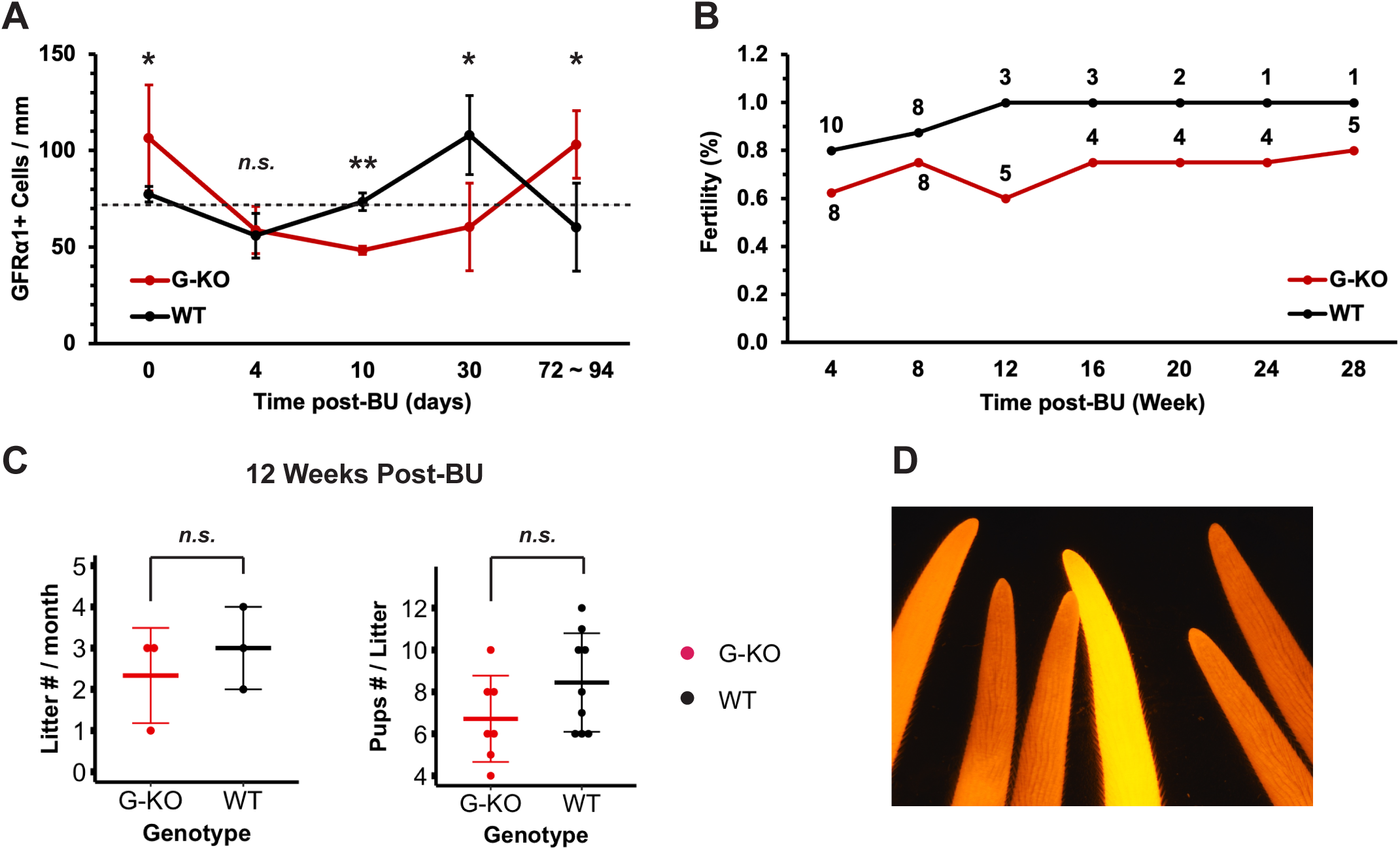
The recoveries of both spermatogonia pool and fertility rate are delayed in adult *Spry4*^G-KO^ males after damage. **(A)** Quantification of GFRα1^+^ cells per mm of seminiferous tubules collected from *Spry4*^G-KO^ or *Spry4*^WT^ males at different time points post-BU. Dashed line, GFRα1^+^ cells per mm in adult *Spry4*^WT^ males at steady state (n>3 mice analyzed per genotype). **(B)** Fertility rate of adult *Spry4*^G-KO^ or *Spry4*^WT^ males post-BU. Number in the linear graph, number of mice tested at the indicated time point. **(C)** Number of litters (left) and offspring pups (right) produced by *Spry4*^G-KO^ or *Spry4*^WT^ males 12 weeks post-BU. **(D)** Red fluorescence (tdTomato) image from the tails of one litter of pups produced by a *Spry4*^G-KO^ male 28 weeks post-BU. Data are mean (SD). *p*-value is calculated by unpaired one-tailed Student’s *t*-test. *n.s.*, not significant (*p*-value >= 0.05).

We also found that although *Spry4*^WT^ mice became completely fertile by 12 weeks post-BU (6-month-old), *Spry4*^G-KO^ mice recovered fertility much slower, with one male remaining sterile within the entire breeding test period (**Figure 5B**). Although the testis-to-body weight ratio remained comparable between *Spry4*^G-KO^ and *Spry4*^WT^ mice across all time points we investigated, *Spry4*^G-KO^ mice exhibited a significantly reduced sperm count compared to *Spry4*^WT^ controls at 26 weeks post-BU, indicating impaired long-term spermatogenic recovery despite normal testicular size (**Figure S13**, and data not shown). All the fertile *Spry4*^G-KO^ males recovered from damage were capable of efficient breeding by producing similar number of litters and offspring pups as *Spry4*^WT^ males (**Figure 5C**). Strong tdTomato signal was observed in every pup produced by *Spry4*^G-KO^ males, revealing efficient GFRα1-Cre^ERT2^-induced recombination within the mature *Spry4*^G-KO^ spermatozoa recovered from regeneration (**Figure 5D**).

### A_undiff_ spermatogonia are progressively reduced in aging *Spry4*^G-KO^ males

We found that aged *Spry4*^G-KO^ mice (> 12-month-old) exhibited a reduction in the overall density of A_undiff_ spermatogonia compared to *Spry4*^WT^ mice at the same age (**Figure S14A**). No significant change in GFRα1^+^ cell density or fraction of GFRα1^+^ clones was observed in *Spry4*^WT^ mice between 4- and 15-months, or between young *Spry4*^G-KO^ and *Spry4*^WT^ mice within 10-months (**Figure S14A** and **S14B**). Interestingly, the reduction in aged mice was accompanied by an enrichment of the most primitive A_s_ spermatogonia (SSCs) population, with a corresponding decrease in A_pr_ and A_al_ clones (**Figure S14C**). These results suggest that, even in the absence of exogenous stress or damage, *Spry4*^G-KO^ spermatogonia progressively adopt a more primitive and less proliferative state with age.

## Discussion

Life-long spermatogenesis requires coordinated self-renewal and balanced differentiation of germ cells to prevent exhaustion of SSCs while maintaining continual germ cell generation. While the molecular and cellular pathways orchestrating spermatogenesis are well documented, the mechanism(s) by which the SSC compartment, which is enriched within a heterogenous population of A_undiff_ spermatogonia capable of dynamic and reversible transition between different developmental phases [3–5], responds to physiological stress to sustain its repertoire of regenerating and differentiating germ cells are less well defined [6, 8]. We have previously shown that MAPK/ERK-dependent negative feedback regulator SPRY4 enables mouse SSCs to respond to growth factor signaling to preserve stem cell activity in culture. Indeed, *Spry4* serves as a marker for adult SSCs in vivo by uniquely expressing in a subset of A_undiff_ spermatogonia but not in differentiating germ cells in adult mouse testis [35]. These observations raised the possibility that SPRY4 might function as a rheostat balancing coordinated germ cell recovery without exhausting the vulnerable SSC compartment. To decipher the cell-autonomous role of SPRY4-ERK signaling in the regeneration of SSCs in vivo, we induced *Spry4* deletion in the spermatogonia of adult mice (*Spry4*^G-KO^), together with a single low-dose BU treatment to deplete most SSCs and rapidly dividing spermatogonia from the testis and followed the recovery of the remaining stem cells over time after injury. We demonstrate that although SPRY4 is dispensable for spermatogenesis in young adult mice, it acts as a critical cell-intrinsic molecular checkpoint of the MAPK/ERK pathway to modulate the progression of balanced germline regeneration, secure rapid germline recovery, and restore long-term fertility in adult mouse testis. Thus, SPRY4 is a novel molecular hub balancing SSC homeostasis under stress to meet the demand for continual generation of mature germ cells.

### SPRY4 modulates SSC responses to genotoxic stress in a phase-specific manner through regulating PI3K/AKT and MAPK/ERK signaling

Based on our study, we propose a model reflecting a phase-specific and SPRY4-dependent adaptation to the regenerative microenvironment following genotoxic stress. Shortly after busulfan-induced damage (BU-D2), the testis is in an acute damage and survival phase when surviving SSCs must prioritize maintenance of an undifferentiated state and stress tolerance. Our data indicate that by poising RTK-dependent MAPK/ERK and PI3K/AKT signaling at basal levels, SPRY4 maintains A_undiff_ spermatogonia (including SSCs) within the wild-type (*Spry4*^WT^) mouse testis in a quiescent state at this stage, thereby preventing surviving A_undiff_ spermatogonia from responding immediately to chemotherapy, limiting premature differentiation, and preserving SSC identity under genotoxic stress. In contrast, BU-D10 represents a distinct biological state with extensive germ cell depletion, coinciding with the onset of regenerative decision-making when surviving SSCs must balance proliferation and differentiation to replenish mature germ cells. We propose that the re-established growth factor signaling, as well as the intercellular interactions between surviving SSCs and the somatic niche, provide transient activation of PI3K/AKT–mTORC1 signaling in SSCs and initiate regenerative proliferation. Under these conditions, MAPK/ERK signaling is also activated and engaged, followed by the induction of its negative feedback regulator SPRY4, which serves to restrain excessive MAPK/ERK activity and prevent premature differentiation, consistent with our prior observation that FGF2 and GDNF induce *Spry4* expression in cultured SSCs [35] (**Figure 6A**).

**Figure 6.**
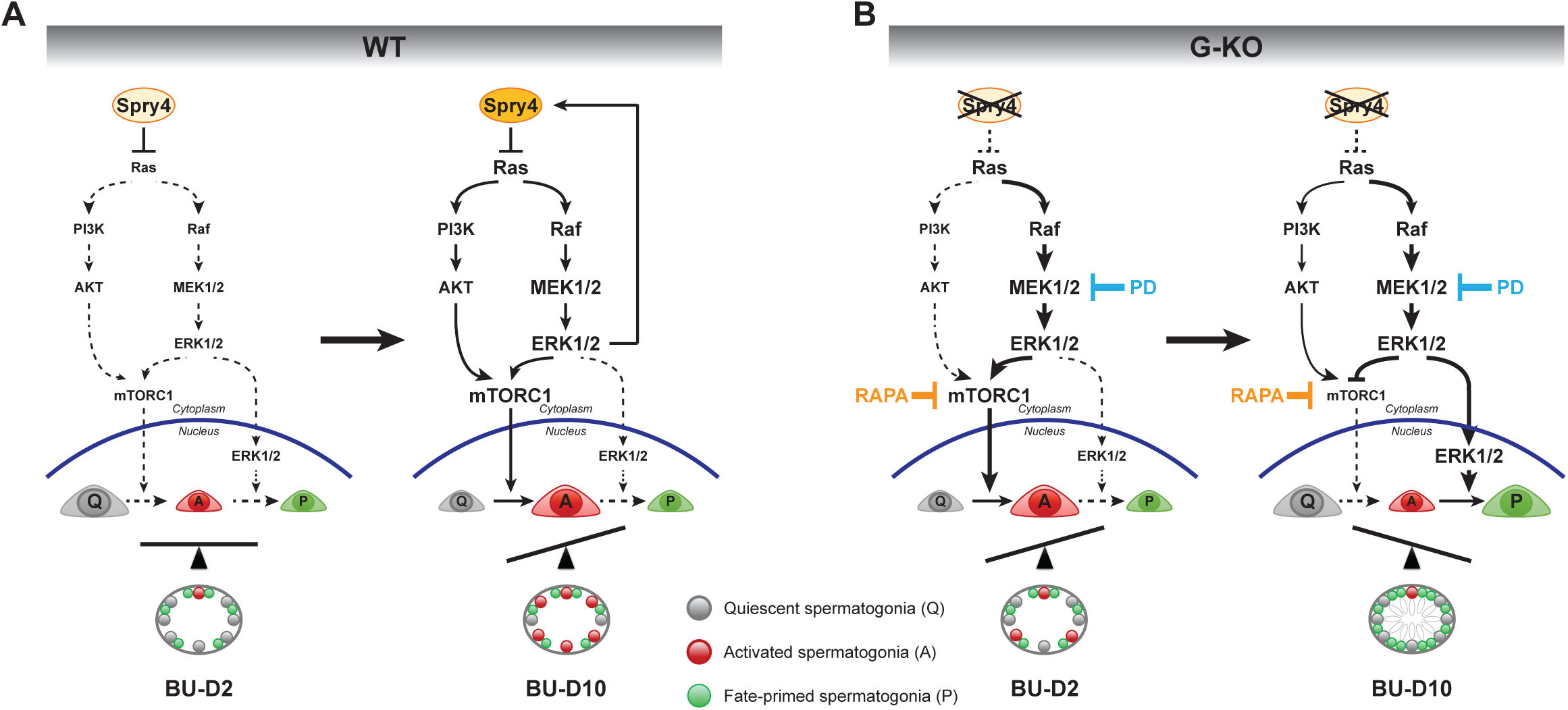
Model illustrating SPRY4-ERK function during spermatogonia recovery and regeneration in BU-damaged mouse testis. **(A)** Wild-type spermatogonia achieve regeneration through a SPRY4-controlled signaling transition. At BU-D2, SPRY4 imposes repression to RTK signaling, inactivating both MAPK/ERK pathway and mTORC1, and poises the surviving A_undiff_ spermatogonia in quiescent state. By BU-D10, with the re-establishment of PI3K/AKT and MAPK/ERK signaling in the recovering A_undiff_ spermatogonia, ERK1/2 is activated in the cytoplasm and cooperates with PI3K/AKT signaling to activate mTORC1. Activated mTORC1 promotes A_undiff_ spermatogonia transition from quiescent to cell-cycle-activated phase (proliferation) and initiates regeneration. Excessive MAPK/ERK signaling and premature differentiation are restrained through reinforcing ERK1/2-dependent SPRY4 activity in A_undiff_ spermatogonia. **(B)** In *Spry4*^G-KO^ spermatogonia, both ERK1/2 and mTORC1 are activated in the cytoplasm of the surviving A_undiff_ spermatogonia immediately after damage (BU-D2), resulting in rapid proliferation in response to injury. By BU-D10, hyperactivated ERK1/2 locates to the nucleus of A_undiff_ spermatogonia and promotes fate commitment, suspending A_undiff_ spermatogonia proliferation with inactivated mTORC1. Q, quiescent A_undiff_ spermatogonia; A, activated A_undiff_ spermatogonia; P, fate-primed spermatogonia; PD, PD0325901; RAPA, Rapamycin.

Deletion of the *Spry4* gene from spermatogonia unleashes RTK-dependent signaling from repression, which activates cytoplasmic ERK1/2 and mTORC1 quickly after damage and induces proliferation in A_undiff_ spermatogonia, resulting in rapid initiation of germ cell regeneration at BU-D2. However, hyperactivated ERK1/2 translocates to the nucleus of A_undiff_ spermatogonia by BU-D10, triggering spermatogonia differentiation while simultaneously blocking mTORC1-supported spermatogonia proliferation (**Figure 6B**). This model is supported by marked enrichment of highly proliferative, differentiation-destined progenitors (RARγ^+^c-KIT^-^) in *Spry4*^G-KO^ testes at an early post-damage time point (BU-D4), whereas the overall A_diff_ spermatogonia (c-KIT^+^) were similarly depleted in both genotypes, indicating that SPRY4 is dispensable to their sensitivity to busulfan. At later stages (BU-D10), the expansion of both progenitors and A_diff_ spermatogonia (MCAM^+^GFRα1^-^) reflects selective amplification of progenitor-derived cells rather than altered initial sensitivity of A_diff_ spermatogonia to chemotoxic injury. Together, these findings indicate that SPRY4 restrains premature proliferative activation of differentiation-destined progenitors during early regeneration, and that the increased progenitors and A_diff_ spermatogonia in *Spry4*^G-KO^ testes at BU-D10 arise from dysregulated regenerative dynamics within A_undiff_ spermatogonia (GFRα1) and progenitors (RARγ), rather than altered busulfan sensitivity.

### The subcellular localization of p-ERK1/2 and pathway cross-regulation collectively determine divergent phase-specific mTORC1 outcomes

ERK1/2 is predominantly retained in the cytoplasm through interactions with anchoring and scaffold proteins in resting cells, whereas phosphorylation-induced conformational changes allow p-ERK1/2 to translocate to other cellular compartments, particularly the nucleus [47]. Cytoplasmic MAPK/ERK signaling preferentially engages PI3K/AKT signaling and supports transient mTORC1 activation, while sustained or excessive nuclear p-ERK1/2 accumulation is more likely to activate transcriptional programs, including differentiation-associated targets and negative feedback regulators that attenuate growth signaling. Consistent with this model, we only observed modest cytoplasmic p-ERK1/2 in *Spry4*^WT^ SSCs during regeneration (BU-D10), with the p-ERK1/2 activity restrained by the ERK negative feedback regulator SPRY4. However, prominent nuclear p-ERK1/2 accumulation in *Spry4*^G-KO^ SSCs at BU-D10 coincided with suppression of mTORC1 activity. Cross-inhibitory interactions between MAPK/ERK and PI3K/AKT pathways may further contribute to the distinct mTORC1 response between *Spry4*^WT^ and *Spry4*^G-KO^ SSCs. Prior studies have shown that excessive MAPK/ERK signaling can negatively regulate PI3K/AKT signaling upstream of mTORC1, i.e., through disruption of adaptor GAB1-mediated PI3K activation [48]. In the absence of SPRY4-mediated feedback restraint, although transient cytoplasmic ERK–mTORC1 coupling was observed in *Spry4*^G-KO^ SSCs shortly after busulfan-induced damage (BU-D2), accumulating hyperactive ERK signaling may progress to dominant inhibition on PI3K/AKT–mTORC1 signaling at a later stage (BU-D10), which can be partially rescued by pharmacologically inhibiting the MAPK/ERK pathway.

### SPRY4-dependent temporal cell-cycle pausing after damage protects SSCs from genotoxic stress and is critical for ensuring appropriate regeneration at a later phase

Studies with multiple model systems, including cancer stem cells (CSCs), reveal that actively cycling cells are particularly vulnerable to alkylating damage, while transient cell-cycle pausing represents a shared strategy by which stem cells survive genotoxic stress. This pause likely limits the propagation of DNA damage and preserves long-term stem cell potential. Our study shows that A_undiff_ spermatogonia undergo a transient regenerative pause following busulfan-induced damage, suggesting that SSC regeneration may represent a physiological counterpart of stress-adaptive mechanisms exploited by CSCs. Importantly, our data suggest that this temporal pause in SSCs is a SPRY4–dependent actively regulated adaptive phase rather than passive quiescence. We found that BU-D2 represents an acute stress-sensing and checkpoint phase, during which surviving A_undiff_ spermatogonia suppress proliferation and differentiation programs while activating stress and damage-response signaling. This pause likely allows stem cells to sense genotoxic stress, assess genomic integrity, and reset signaling thresholds (i.e., mTORC1, ERK1/2) before entering the regenerative phase at BU-D10. Unleashing the checkpoint at BU-D2 is detrimental to stem cells, and the SPRY4–ERK signaling appears to play a key role in controlling the timing of exit from this paused state, thereby licensing appropriate regenerative responses. In the absence of SPRY4, this timing becomes deregulated, resulting in premature differentiation and compromised regeneration, an outcome reminiscent of dysregulated stress responses observed in CSCs.

### SPRY-ERK signaling modulates stress tolerance thresholds in SSCs and impacts surveillance under stress

Loss of *Spry4* leads to persistent accumulation of DNA damage, as marked by sustained γH2A.X enrichment, in both primitive SSCs (GFRα1^+^PLVAP^+^) and differentiation-primed A_undiff_ spermatogonia (GFRα1^+^PLVAP^-^) following genotoxic injury. Notably, this elevated DNA damage persists from early regeneration (BU-D10) through later recovery phases (BU-D30), when wild-type spermatogonia have largely resolved stress responses and resumed steady-state proliferation. Although *Spry4*^G-KO^ testes retain A_s_ spermatogonia expressing primitive SSC markers, a substantial fraction of these cells exhibit prolonged genotoxic stress, indicating a compromised cellular state rather than a fully functional, homeostatic SSC pool. Consistent with this interpretation, *Spry4* deletion delays recovery of A_undiff_ spermatogonia after injury, reduces long-term sperm output and fertility, and is associated with progressive adoption of a more primitive and hypoproliferative state with age even in the absence of exogenous damage. Together, these data suggest that SPRY4–ERK signaling contributes to SSC stress tolerance and recovery, potentially by interfacing with genome integrity surveillance pathways that regulate DNA repair capacity or damage tolerance thresholds. While persistent γH2A.X signaling raises the possibility that a subset of SSCs may enter senescent or senescence-like states, technical limitations precluded direct assessment of canonical senescence markers in this study. Future work incorporating optimized detection of p21, p16, and related pathways, as well as analysis of ATM–Trp53 signaling, will be necessary to define how SPRY4–ERK signaling mechanistically integrates genome integrity control with long-term SSC function.

### SPRY4 functions as a context-dependent regulator of MAPK/ERK signaling in SSCs: in vivo versus in vitro

Previous studies, including foundational work from the Shinohara group and our own analyses in cultured SSCs, established Ras/Raf/MEK/ERK signaling cascade downstream of GDNF and FGF2 as a central driver of SSC proliferation and self-renewal under steady-state or growth factor–rich in vitro conditions [23, 24, 35, 49]. Those studies identified ERK-responsive transcriptional programs involving immediate-early genes (*c-FOS*), cell-cycle regulators (cyclins A/D/E, CDK2), and ERK-dependent phosphorylation of transcription factors (CREB-1, ATF-1, and CREM-1). In our prior in vitro work [35], we demonstrated that SPRY4 functions as an ERK-induced negative-feedback regulator required to preserve SSC stemness by constraining MAPK/ERK signaling amplitude; loss of SPRY4 in culture led to ERK dysregulation, premature differentiation, and reduced transplantation capacity.

The present study extends these findings to a physiologically relevant in vivo regeneration model, revealing that SPRY4 is not uniformly required for SSC maintenance under homeostasis but becomes functionally essential under regenerative stress following genotoxic injury. Germline-specific *Spry4* deletion leads to ERK hyperactivation, premature differentiation, delayed recovery of A_undiff_ spermatogonia, and compromised long-term fertility. Importantly, this context dependence reconciles apparent differences between in vitro and in vivo phenotypes: while cultured SSCs are highly dependent on SPRY4-mediated feedback under growth factor–rich conditions, SSCs in vivo can partially compensate for SPRY4 loss at steady state, likely through redundant feedback regulators (e.g., SPRY1/2, DUSPs). Our findings therefore resolve the apparent discrepancy between in vitro and in vivo phenotypes and establish SPRY4 as a stress-revealed MAPK/ERK signaling rheostat, required to coordinate SSC fate decisions specifically during regenerative challenge, when the precise temporal and spatial control of ERK signaling by SPRY4 is particularly critical for SSCs to balance stress adaptation, proliferation, and differentiation.

### SPRY4–ERK target genes in regenerative SSCs: extending beyond proliferative outputs

In contrast to prior studies that primarily linked ERK signaling to proliferation-associated transcriptional programs, our study identifies several novel SPRY4–ERK–responsive genes during SSC regeneration (e.g., *Id1* and *Cxcl12*), highlighting a distinct class of downstream targets engaged under genotoxic stress. Notably, expression of both *Id1* and *Cxcl12* was restricted to specific spermatogonial progenitor subsets rather than broadly induced across the A_undiff_ spermatogonia, indicating a stage- and context-dependent transcriptional response.

ID proteins are well established as regulators of stem cell fate in multiple tissues, where they preserve stemness by antagonizing E-protein–mediated differentiation programs [39]. Although ID4 has been implicated in SSC self-renewal, the function of ID1 in mammalian SSCs has remained undefined [50–53]. Our data suggest that SPRY4–ERK–dependent regulation of *Id1* supports SSC regenerative competence, rather than steady-state self-renewal, as *Id1* expression is reduced in *Spry4*-deficient spermatogonia during regeneration and restored upon MAPK/ERK pathway inhibition. These findings raise the possibility that ID proteins act as key executors of SPRY4-mediated ERK feedback during SSC regeneration. Similarly, while CXCL12 has been studied as a niche-derived chemokine critical for SSC homing and maintenance in transplantation assays, its SSC-intrinsic role has not been directly demonstrated [43, 44]. Our scRNA-seq analysis indicates that *Cxcl12* expression is confined to a defined progenitor subset in the adult testis and is dynamically regulated by SPRY4–ERK signaling during regeneration. This supports a context-dependent, signaling-responsive role for CXCL12, rather than an obligatory function in SSC maintenance.

### Regulation of SPRY4 beyond canonical negative feedback from MAPK/ERK signaling

As one of the major negative-feedback regulators of RTK-dependent signaling pathways, SPRY4 activity can be tightly controlled through its intracellular localization, post-translational modification, or binding with cofactors including other Sprouty family isoforms [54]. Our study in wild-type A_undiff_ spermatogonia revealed that SSCs activate their regenerative responses through RTK-dependent PI3K/AKT and MAPK/ERK pathways at least two days after chemotherapy-induced damage. Deletion of the *Spry4* gene in spermatogonia augmented SSC regeneration. Given that *Spry4* is exclusively expressed in A_undiff_ spermatogonia within the adult mouse germ cells [35], which presumably inhibits RTK-dependent signaling pathways in SSCs at steady state and immediately after injury (within 2 days of BU treatment), pausing SPRY4 activity in SSCs would be required to initiate regeneration in wild-type mice.

It has been reported that a hetero-oligomer formed between SPRY4 and SPRY1 exhibits the most potent inhibitory effect on ERK activation in cultured mouse or human cells [55], suggesting that the activities of other Sprouty family isoforms in A_undiff_ spermatogonia may also facilitate SPRY4 function during regeneration. Our study identified the role of SSC-intrinsic SPRY4 in responding to damage, while further investigation about how to regulate the activities of SPRY4 and other Sprouty family isoforms in germ cells will be needed. Considering the essential role of the stem cell microenvironment in maintaining and regulating stem cell function, somatic cells that constitute SSC niche (i.e., Sertoli, Leydig, peritubular myoid and testicular endothelial cells) may also modulate SPRY4 activities within SSCs and promote regeneration after injury through multiple mechanisms. Indeed, Sertoli cells have been reported to periodically activate ERK1/2 and produce GDNF to support SSC self-renewal at selected stage of seminiferous cycle [26]. Specialized testicular endothelial cells have also been identified as a critical population in the germline stem cell niche by producing GDNF and other factors to support human and mouse SSCs in culture [56].

Both somatic stem cells and SSCs are vulnerable to DNA damage induced by chemotherapy, resulting in severe side effects or long-term tissue dysfunction (i.e. infertility) in cancer patients post-treatment [57]. Specifically, there are no means to preserve the fertility of prepubertal male patients, who are not yet producing mature sperms and are therefore unable to benefit from standard sperm banking [58]. Elucidation of the signaling pathways and regulatory networks that control stem cell self-renewal, proliferation, and differentiation in adolescent and adult organs is essential to develop targeted therapies, improve tissue repair, and advance regenerative medicine. By dissecting the key transition steps in recovering SSCs, we identified SPRY4-ERK signaling as the first critical checkpoint regulator that modulates germline-specific damage response to restore male reproduction in adult mammals. Our study sets forth the notion that after chemotherapy acutely depletes testicular cells, particularly the rapidly dividing spermatogonia, and disrupts intercellular signals required for SSC homeostasis, the surviving stem cells rely on SPRY4-ERK signaling to execute a phase-specific regeneration program and ensure appropriate germline recovery without exhausting SSCs. During the progression of regeneration, SPRY4 is required to appropriately balance stem cell proliferation with differentiation through regulating ERK1/2 activity. Loss of *Spry4* in A_undiff_ spermatogonia accelerates the regeneration response to BU damage, with hyperactivated ERK1/2 further promoting spermatogonia differentiation at the expense of self-renewal. Furthermore, chronological aging appears to push the SPRY4-deficient SSC pool toward a more quiescent, primitive state, potentially compromising the long-term regenerative capacity of the germline, providing further evidence that SPRY4 plays a key role in maintaining the homeostatic balance between SSC self-renewal and differentiation in steady-state mice without overt damage. Our results reveal a novel negative feedback regulator-dominant mechanism in orchestrating cell-intrinsic signaling to coordinate adult stem cell recovery and long-term homeostasis, shedding a light on improving tissue regeneration after drug induced injury and preserving human fertility from environmental stresses.

## Materials and methods

### Ethics statement

This study was approved by the Weill Cornell Medical College IACUC (#2010-0028). For euthanasia, mice were exposed to CO_2_ followed by cervical dislocation.

### Mouse maintenance and treatment

The *Gfr*α*1-CreER^T2^*mice were a gift from Dr. Sanjay Jain [4, 59]. *Rosa26-lox-stop-lox(LSL)-tdTomato* reporter mice were obtained from Jackson Laboratory (strain #: 007914) [60]. The *Spry4^flox^* mice were a gift from Dr. Ophir Klein [61]. The genotype of the newborn mice was determined by PCR with the primers listed below. For gene deletion, tamoxifen (HY-13757A, MedChemExpress) was administered to adult mice (2- to 4-month-old) by intraperitoneal (IP) injection at a dose of 100 mg/kg of body weight once every 24 h for 4 consecutive days per week for 2 weeks total. To induce regeneration, busulfan (HY-B0245, MedChemExpress) was administered to mice by IP injection at a single dose of 10 mg/kg of body weight. BU-treated cohorts were treated daily by IP injection starting from 3 days post-BU for 7 consecutive days with PD0325901 (HY-10254, MedChemExpress, 2.5 mg/kg), Rapamycin (HY-10219, MedChemExpress, 4 mg/kg) or vehicle (10% DMSO/PBS). The volume of drug or vehicle was adjusted based on mouse body weight. No overt toxicity (e.g., weight loss, lethargy, or histological damage) was observed in vehicle-treated mice.

### Sperm counting

After euthanizing the mice, one cauda epididymide was dissected from each mouse and thoroughly minced in a 1.5 ml microcentrifuge tube containing 1 ml PBS. The sample was left in room temperature for 15 min to allow the sperm to swim out. The sperm suspension was diluted 10 folds with PBS and counted on a hemocytometer. Sperm counts were obtained twice and recorded as the average of the two counts.

### Hematoxylin and eosin (H&E)

Tissues were fixed with 4% paraformaldehyde/PBS overnight at 4 □C, washed and immersed in PBS. H&E staining was performed at center for translational pathology at Weill Cornell Medicine following standard methods. Images were captured with an ECHO Revolve microscope and tissue architecture was scored blindly.

### Whole-mount immunofluorescence (IF)

For whole-mount IF, seminiferous tubules were teased apart from freshly collected detunicated testes and rinsed in PBS. Tubules were fixed with 4% PFA (158127, Sigma-Aldrich) overnight at 4°C and then washed with PBS for three times. The tubules were incubated with the basic optical clearing solution consists of 150mM KCl, 20% DMSO, 0.5% triton, 0.5% NaN3, and 10mM Tris-HCl (pH 9.5) for 20 min at 37°C to decolorize the tissue and reduce autofluorescence [62]. Tubules were then blocked with 1 % normal donkey serum albumin (A1470,Sigma-Aldrich) for 1 hr at room temperature in PBS before incubation with 1 μg/ml goat anti-GFRα1 (AF560, R&D Systems), 1 μg/ml rat anti-MCAM antibody (134702, BioLegend), 1:250 rabbit anti-phospho-p44/42 MAPK (Erk1/2) antibody (4370S, Cell Signaling), 1 μg/ml Ki67 (ab15580, Abcam), or 1:500 rabbit anti-pRPS6 (9101S, Cell Signaling), overnight at 4°C in a nutating mixer at 4°C. Detection of primary antibodies were performed using 1 μg/ml Alexa-647 conjugated Anti-Rabbit IgG (711-065-152, Jackson ImmunoResearch), Alexa-488 conjugated Anti-Goat IgG (705-545-147, Jackson ImmunoResearch), anti-rat IgG 568 (A-11077, Invitrogen), or anti-rabbit IgG 647 (711-605-152, Jackson ImmunoResearch) for 60 min at room temperature. Note, all the incubation were done in the nutating mixer to ensure proper penetration of solutions. The tubules were mounted in PBS with DAPI in a 35-mm glass-bottomed dish (Part No: P35G-0-14-C, MatTek) and top with a glass cover slip. Image analysis was performed on a Zeiss LSM800 confocal microscope.

### IF image analysis

To quantify spermatogonia in adult mouse testis, the seminiferous tubules were examined by whole-mount IF staining for GFRα1. The density and the fraction of A_s_, A_pr_, and A_al_ spermatogonia were calculated by scoring n>100 GFRα1^+^ cells within over 5 mm of seminiferous tubules per animal. 3 – 5 animals were analyzed per condition.

### Fluorescence-activated cell sorting (FACS)

To isolate spermatogonia, testes were collected from adult mice 10 days after recovering from BU treatment (BU-D10). Single-cell suspensions were generated from testes by enzymatic digestion as described before [63]. Briefly, seminiferous tubules were collected from detunicated testes and minced on ice. The tissue was enzymatically dissociated with agitation for 30 min at 37°C in a buffer containing 0.017% trypsin (Cellgro), 17mM EDTA (Cellgro), 0.03% collagenase (Sigma-Aldrich) and DNase I (100 μg/ml; Roche). For fluorescence-activated cell sorting (FACS), spermatogonia were firstly enriched for MCAM/CD146^+^, a marker of SSCs and spermatogonia [36, 37], via mouse CD146 (LSEC) MicroBeads (Miltenyi Biotec, 130-092-007), then proceeded with antibody staining with Alexa Fluor 647 anti-MCAM antibody (clone ME-9F1, 134718, BioLegend), Alexa Fluor 488 anti-c-KIT (CD117) antibody (clone 2B8, 105816, BioLegend), and PE/Cyanine7 anti-CD31 antibody (clone WM59, 303118, BioLegend) each at 10 μg/ml for 30 min at 4°C. After bead enrichment, about 1 – 2 x10^6^ testicular cells were collected for FACS. Cell suspensions with immunocomplexes were subject to FACS with a BD FACS Aria II instrument (Beckman Coulter, USA). DAPI (D1306, Thermo Fisher Scientific) was used for live/dead cell discrimination. After exclusion of doublets and DAPI-positive cells, as well as testicular endothelial cells (CD31^+^), the undifferentiated germ cell population (MCAM^High^c-KIT^-^, including GFRα1^+^ A_undiff_ spermatogonia) were gated and collected.

### FACS analysis

FACS results were analyzed with Flowjo 9.3.2. <u>S</u>SC <u>D</u>ifferentiation Index (SDI) was calculated by: SDI = Fraction of MCAM^Low^CD31^-^ cells / Fraction of MCAM^High^c-KIT^-^CD31^-^ cells

### RNA extraction

Total RNA was extracted from fresh spermatogonia isolated by FACS using PicoPure RNA isolation kit (KIT0204, Applied Biosystems) according to the manufacturer’s instructions. Residual genomic DNA was removed by on-column DNA digestion (79254, Qiagen).

### Library construction and sequencing

The stranded mRNA sequence libraries were generated using the KAPA mRNA HyperPrep Kit (Kapa Biosystems, KK8580) according to the manufacturer’s instructions. Sequencing service was performed on an Illumina NovaSeq X plus sequencer according to the standard Illumina protocol.

### Quantitative reverse transcription polymerase chain reaction (qRT-PCR)

qRT-PCR was performed in triplicate in a LightCycler 480II Real-Time System (Roche Molecular Systems) using Luna Universal qPCR Master Mix (M3003E, NEB). Primers were validated for use based on efficiency and inspection of melting curves. For quantification, each technical triplicate was normalized to endogenous *Tbp* and relative transcript expression was calculated using the comparative C_T_ method (2^−ΔΔ^*Ct* method) using the average value of the technical triplicates of the control condition as the reference value. Data presented correspond to *Tbp* normalization. The qRT-PCR primers are listed below.

### RNA-seq data processing and analysis

Sequencing reads were de-multiplexed, checked for quality (FastQC), and trimmed/filtered when appropriate. The resultant high quality reads were mapped (STAR v2.7.10a) to the transcriptome sequence reference of the GENCODE mm10 GRCm39 version 107 build. Gene counts and transcript abundance measures (FPKM values) were quantified using the R package edgeR (v4.0.14). Gene expression was reported as log2 transformed FPKM value. With Limma R package and expression profiles generated in our labs, lists of the differentially expressed genes in the unpair-wise comparisons between gene deletion and control samples were generated with at least one-fold (log2) change and statistical significances (*p*-value < 0.05). At least three biological replicates from each sample were analyzed.

### Statistical analyses

Results are presented as means ± SD. At least three biological replicates and three technical replicates were performed for each experiment unless otherwise indicated in the text. All comparisons were performed using an unpaired one-tailed Student’s *t*-test unless otherwise noted. Results were considered significant at *p*-value <0.05. R (version 4.4.1) was used for statistical analyses and generating graphs.

### Primer sequences for genotyping and qRT-PCR

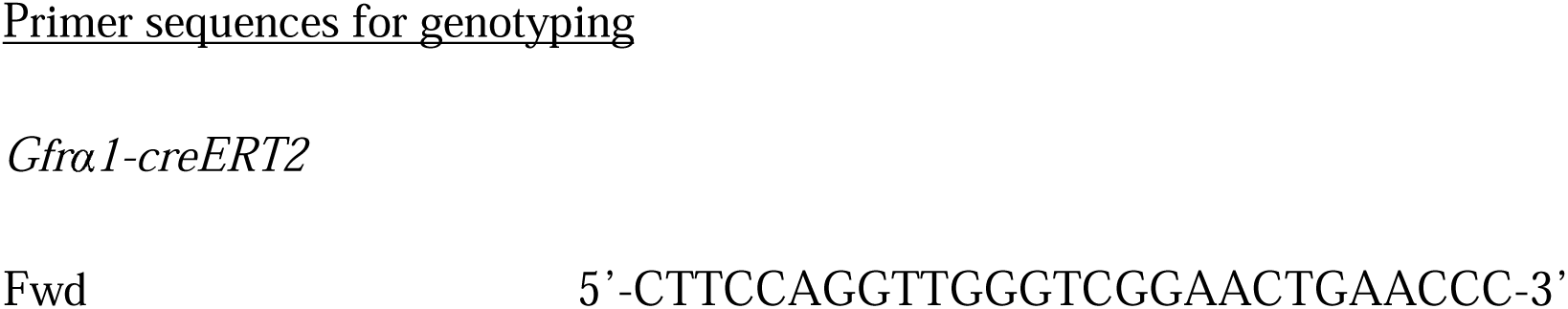
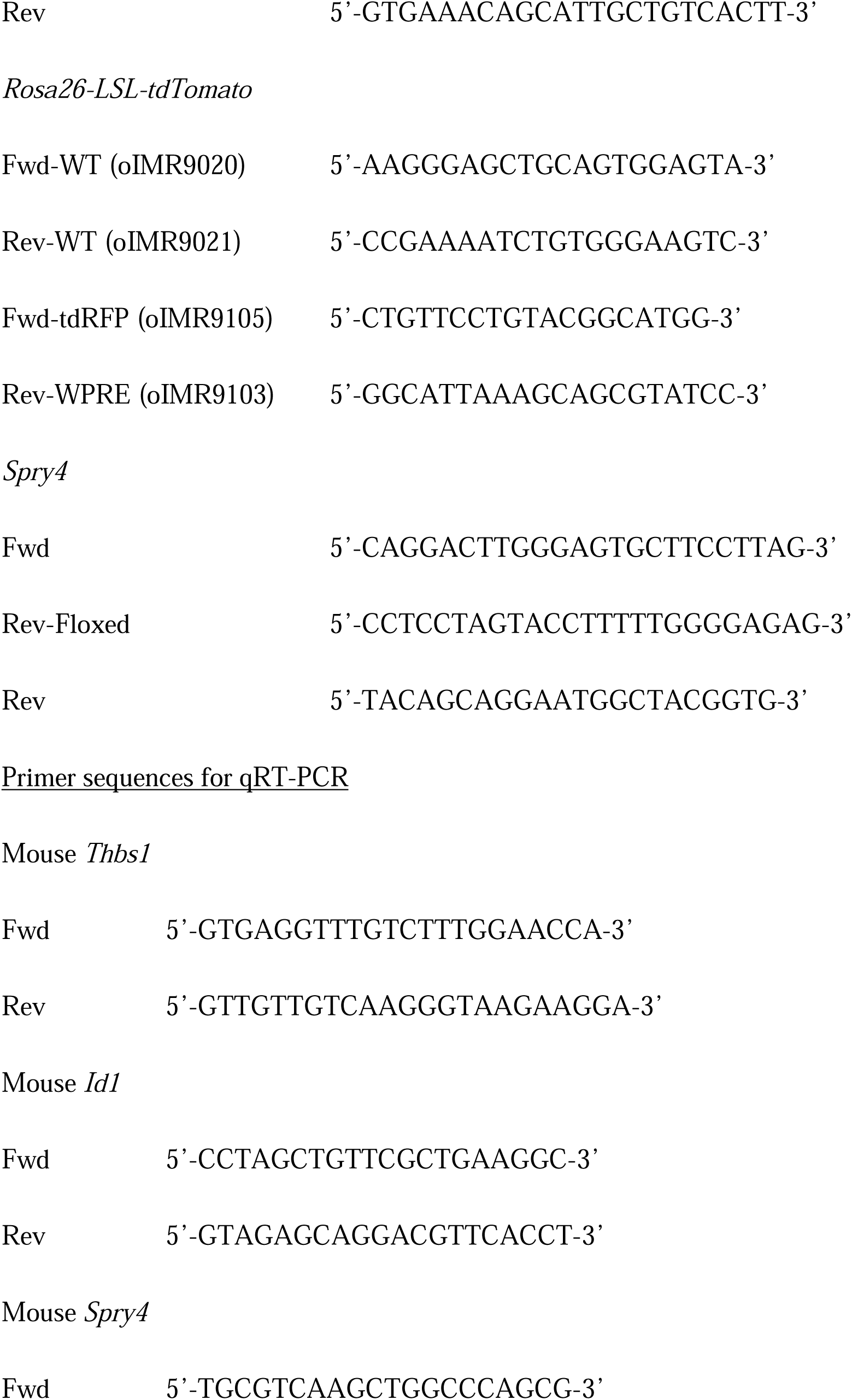
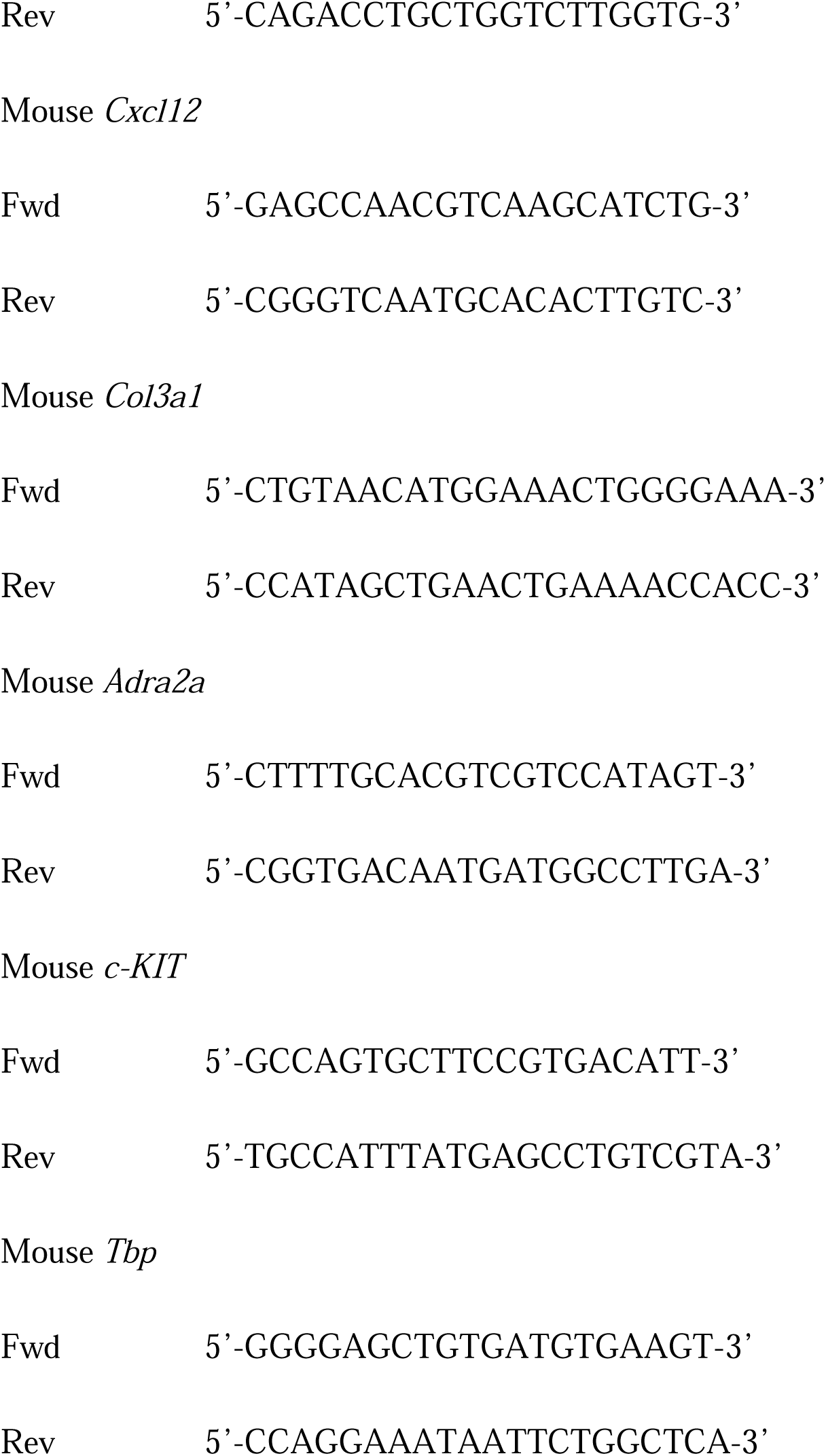

## Acknowledgments

This work was supported by grants from the National Institutes of Health (R01HD101452 and R21HD107212 to T.E. and Y.L.) and LaGuardia Community College Bridges to the Baccalaureate Program to (W.W. and N.X.). We thank Dr. Jenny Xiang and the Genomics Resources Core Facility at Weill Cornell Medicine for performing RNA-seq analysis. We thank the Flow Cytometry Core Facility at Weill Cornell Medicine for their timely help and discussions on flow cytometry analysis and FACS. We thank Henry Mejia and Jiale Lin at LaGuardia Community College for helping with mouse genotyping and organ collection. We thank Dr. Lonny R. Levin, Dr. Jochen Buck, and Dr. Justine Fischoeder for insightful discussions and intellectual input during the course of this work. We thank Odin Wright at LaGuardia Community College for consultation in the bioinformatics analyses. All authors declare that they have no conflicts of interest.

## Data Availability Statements

The data underlying this article are available in the article, in its online supplementary material, and at the following link: (https://tbd).

## Supplementary Data

**Supplementary Figure 1. Fluorescent reporter in *Spry4*^G-KO^ testis labels germ cells of all stages.** Testes collected from adult *Spry4*^G-KO^ (G-KO) mice 6 weeks post-tamoxifen treatment were analyzed by fluorescent microscopy for **(A)** tdTomato (tdT) expression in whole-mount seminiferous tubules. Red, tdTomato; Blue, DAPI. Representative whole-mount IF images illustrating tdTomato^+^ populations (Red) in tubules collected from *Spry4*^G-KO^ mice. **(B)** Light blue, VASA; **(C)** Magenta, GFRα1; Green, MCAM; **(D)** Grey, Vimentin; Green, MCAM; Blue, DAPI. Inset, higher magnification images of indicated regions without DAPI or tdT.

**Supplementary Figure 2. Characterization of germline-specific *Spry4* deletion in adult mouse testis at steady state.** Testes collected from adult (within 4 months of age) *Spry4*^G-KO^ (G-KO) or *Spry4*^WT^ (WT) mice 6 weeks post-tamoxifen treatment were compared for **(A)** testis to body weight ratio and **(B)** sperm count. Data are mean (SD), *p*-value is calculated by unpaired one-tailed Student’s *t*-test. *n.s.*, not significant (*p*-value >= 0.05). n>5 mice analyzed per genotype. **(C)** H&E–stained histological cross-sections of testis (up), caput and cauda epididymis (down).

**Supplementary Figure 3. H&E–stained histological cross-sections of mouse testes and epididymides.** Organs collected from adult (4-month-old) *Spry4*^G-KO^ (G-KO) or *Spry4*^WT^ (WT) mice treated with **(A)** single low-dose BU (10 mg/kg) or **(B)** DMSO. Testis (up) and epididymis (down) were collected 10 days after treatment. These images reveal the lower magnification areas from **Fig 1D**.

**Supplementary Figure 4. Characterization of A_undiff_ spermatogonia with PLVAP. (A)** Whole-mount IF staining of tubules collected from adult *Spry4*^G-KO^ (G-KO) or *Spry4*^WT^ (WT) mice at BU-D10. Inset, higher magnification images of indicated regions. Magenta, GFRα1; Green, PLVAP; Blue, DAPI. Arrowhead, GFRα1^+^PLVAP^+^ A_s_ spermatogonia; Asterisk, GFRα1^+^PLVAP^-^ A_s_ spermatogonia; Brackets, A_pr_ spermatogonia; Dashed outline, A_al_ spermatogonia (4 – 8 cells). **(B)** Quantification of PLVAP^+^ cells within A_s_ spermatogonia (GFRα1^+^) in whole-mount IF (n>3 mice analyzed per condition, n>50 GFRα1^+^ cells scored per animal). *n.s.*, not significant (*p*-value >= 0.05). Data are mean (SD), *p*-value is calculated by unpaired one-tailed Student’s *t*-test.

**Supplementary Figure 5. Flow cytometry analysis of MCAM^+^ spermatogonia at BU-D10.** Testicular cells collected from adult mice at BU-D10 were analyzed after exclusion of doublets, dead cells (DAPI^+^), and testicular endothelial cells (CD31^+^). **(A)** Representative flow cytometry of MCAM^+^ testicular cells for Ki67. Red, *Spry4*^G-KO^ (G-KO); Blue, *Spry4*^WT^ (WT); Grey, negative control with unstained cells. **(B)** Percentage of cells within Ki67^+^ gates. n=3 mice analyzed per condition. Data are mean (SD), *p*-value is calculated by unpaired one-tailed Student’s *t*-test.

**Supplementary Figure 6. Flow cytometry analysis of A_diff_ spermatogonia at BU-D4.** Testicular cells collected from adult mice at BU-D4 were analyzed after exclusion of doublets, dead cells (DAPI^+^), and testicular endothelial cells (CD31^+^). **(A)** Marker expression associated with distinct populations of spermatogonia. P, progenitors; D, A_diff_ spermatogonia. **(B)** Representative flow cytometry of testicular cells for RARγ and c-KIT. Percentages of cells within RARγ^+^c-KIT^-^ gates are indicated. **(C)** Percentage of cells within RARγ^+^c-KIT^-^ gates. **(D)** Representative flow cytometry analysis for c-KIT. **(E)** Percentage of cells within c-KIT^+^ gates. **(F)** Representative flow cytometry analysis for Ki67 within RARγ^+^ cells. **(G)** Percentage of Ki67^+^ cells within RARγ^+^ cells. n>3 mice analyzed per condition. *n.s.*, not significant (*p*- value >= 0.05). Data are mean (SD), *p*-value is calculated by unpaired one-tailed Student’s *t*-test.

**Supplementary Figure 7. FACS strategy for isolating spermatogonia.** Testicular cells collected from adult mice at BU-D10 were first enriched for MCAM/CD146 expression by MicroBeads, then FACS-isolated for spermatogonia after **(A)** exclusion of doublets, dead cells (DAPI^+^), and testicular endothelial cells (CD31^+^). **(B)** Selected testicular cells were gated for MCAM^+^ and c-KIT^-^ populations based on single antibody staining. **(C)** This strategy enabled the isolation of two populations, A_undiff_ spermatogonia (MCAM^High^c-KIT^-^CD31^-^) and A_diff_ spermatogonia (MCAM^Low^CD31^-^), from mouse CD146 MicroBeads-enriched testicular cells. **(D)** tdTomato^+^ cells within MCAM^High^c-KIT^-^ cells isolated from *Spry4*^G-KO^ mouse testes (Red). Grey, MCAM^High^c-KIT^-^ cells isolated from *Spry4*^WT^ mouse testes that did not express tdTomato reporter.

**Supplementary Figure 8. Characterization of A_undiff_ spermatogonia with PD0325901 (PD) treatment.** Seminiferous tubules collected at BU-D10 after BU+PD (PD) or BU+DMSO (DMSO) treatment were analyzed by whole-mount IF staining for GFRα1 expression, and quantified for GFRα1^+^ cells per mm of tubules in **(A)** *Spry4*^WT^ (WT) or **(B)** *Spry4*^G-KO^ (G-KO) mice, or fraction of A_s_, A_pr_, and A_al_ spermatogonia within the GFRα1^+^ clones in **(C)** WT or **(D)** G-KO mice. Numbers in round brackets indicate cell numbers per GFRα1^+^ clone (n=3 mice analyzed per genotype). Data are mean (SD). *p*-value is calculated by unpaired one-tailed Student’s *t*-test. *n.s.*, not significant (*p*-value >= 0.05).

**Supplementary Figure 9. Expression of selected genes in vivo. (A)** qRT-PCR analysis of selected genes. Data are mean (SD). *p*-value is calculated by unpaired one-tailed Student’s *t*-test. *, *p*-value < 0.05; **, *p*-value < 0.01. **(B)** Expression of selected genes in each spermatogonia subpopulation in vivo. The single-cell RNA sequencing data was collected from Tomato^+^Venus^+^ (Spry4^+^) germ cells FACS-isolated from tamoxifen-induced GCRT;*Spry4*^H2B-Venus^ mice and analyzed by UMAP clustering to define five major populations, including early and late SSCs that highly express SSC markers, and progenitors Spg1 (1) / Spg2 (2) / Spg3 (3) that exhibit low expression of SSC markers, as reported previously [35]. **(C)** Cxcl12 expression (left) within a subset of progenitors (mainly Spg2) compared to UMAP clustering from all Spry4^+^ spermatogonia (right, adapted from Luo Y, et al., *Biol. Reprod.* 2023).

**Supplementary Figure 10. ERK activity within the seminiferous tubules during regeneration (BU-D10).** Adult mice were treated with a single low-dose BU and then daily with DMSO (BU+DMSO), PD0325901 (BU+PD), or Rapamycin (BU+RAPA) starting from day 3 post-BU. Testes were collected for analysis at BU-D10 by whole-mount IF staining for ERK activity in the seminiferous tubules collected from **(A)** *Spry4*^WT^ mice. Red, MCAM; Cyan, p-ERK1/2; Blue, DAPI, or **(B)** *Spry4*^G-KO^ mice. Red, tdTomato; Grey, GFRα1; Cyan, p-ERK1/2; Blue, DAPI. Inset, higher magnification images of indicated regions without DAPI. Asterisks, somatic cells with p-ERK1/2.

**Supplementary Figure 11. ERK activity within the seminiferous tubules quickly after damage (BU-D2).** Adult mice were treated with a single low-dose BU and then daily with DMSO (BU+DMSO), PD0325901 (BU+PD), or Rapamycin (BU+RAPA) starting from day 0 post-BU. Testes were collected for analysis at BU-D2 by whole-mount IF staining for ERK activity in the seminiferous tubules collected from **(A)** *Spry4*^WT^ mice. Red, MCAM; Cyan, p-ERK1/2; Blue, DAPI, or **(B)** *Spry4*^G-KO^ mice. Red, tdTomato; Grey, GFRα1; Cyan, p-ERK1/2; Blue, DAPI. Inset, higher magnification images of indicated regions without DAPI. Asterisks, somatic cells with p-ERK1/2.

**Supplementary Figure 12. Spermatogonia remain reduced in adult *Spry4*^G-KO^ mice 30 days after damage (BU-D30). (A)** Testes collected from adult G-KO or WT mice at 30 days post-BU were analyzed by whole-mount IF for GFRα1^+^ populations. Higher magnification images of indicated regions are shown on the right. Magenta, GFRα1; Blue, DAPI. Asterisks, A_s_ spermatogonia; Brackets, A_pr_ spermatogonia; Dashed lines, A_al_ spermatogonia (4 – 8 cells). **(B)** Quantification of GFRα1^+^ cells per mm of seminiferous tubules. **(C)** Fraction of A_s_, A_pr_, and A_al_ spermatogonia based on whole-mount IF analysis of GFRα1^+^ clones. Number in round brackets indicates cell numbers per GFRα1^+^ clone. **(D)** Whole-mount IF staining on γH2A.X foci within the nuclei of A_undiff_ spermatogonia collected at BU-D30. Higher magnification images of the indicated regions are shown below. Magenta, GFRα1; Green, PLVAP; Grey, γH2A.X; Blue, DAPI. Arrowhead, GFRα1^+^PLVAP^+^ A_s_ spermatogonia; Asterisk, GFRα1^+^PLVAP^-^ A_s_ spermatogonia; Dashed outline, GFRα1^+^ A_s_ spermatogonia. **(E)** Quantification of γH2A.X^+^ cells within A_s_ spermatogonia (GFRα1^+^) in whole-mount IF staining of tubules collected at BU-D10 (pink) or BU-D30 (dark red). Dashed line, fraction of γH2A.X^+^ cells within GFRα1^+^ cells at steady state in wild-type mouse testis. (n>100 GFRα1^+^ cells scored per animal). Data are mean (SD), n>3 mice analyzed per genotype. *p*-value is calculated by unpaired one-tailed Student’s *t*-test.

**Supplementary Figure 13. Characterization of adult mouse testes recovered long-term after damage.** Testes collected from adult *Spry4*^G-KO^ (G-KO) or *Spry4*^WT^ (WT) mice 26 weeks post-busulfan treatment were compared for **(A)** testis to body weight ratio and **(B)** sperm count. *n.s.*, not significant (*p*-value >= 0.05). Data are mean (SD), *p*-value is calculated by unpaired one-tailed Student’s *t*-test. n>3 mice analyzed per genotype.

**Supplementary Figure 14. A_undiff_ spermatogonia are progressively reduced and enriched to A_s_ spermatogonia in aging *Spry4*^G-KO^ males.** Testes collected from adult *Spry4*^G-KO^ or *Spry4*^WT^ mice at different ages were analyzed by whole-mount IF for GFRα1^+^ populations. **(A)** Quantification of GFRα1^+^ cells per mm of seminiferous tubules. Based on whole-mount IF analysis of GFRα1^+^ clones, the fractions of A_s_, A_pr_, and A_al_ spermatogonia were quantified for **(B)** young mice (4- ∼ 5-month-old, 6 weeks post-BU) and **(C)** old mice (13- ∼ 15-month-old, 42- ∼ 48-week post-BU). Number in round brackets indicates cell numbers per GFRα1^+^ clone. Data are mean (SD), n>3 mice analyzed per genotype. *p*-value is calculated by unpaired one-tailed Student’s *t*-test.

## Notes

### Competing Interest Statement

The authors have declared no competing interest.

### Summary of Updates

Introduction revised to clarify current models of spermatogonial stem cell plasticity and to better position the study within the context of regenerative spermatogenesis and stress response signaling. Results section updated to include new analyses demonstrating increased DNA damage and impaired genome integrity in Spry4 deficient undifferentiated spermatogonia during regeneration, including quantification of γH2A.X foci after busulfan treatment; Section on early regenerative dynamics expanded to clarify the temporal sequence of proliferation and differentiation defects in Spry4 knockout testes at multiple recovery time points. Flow cytometry and whole-mount IF analyses revised to improve characterization of spermatogonial subpopulations, including PLVAP+ stem cells, RARγ+ progenitors, and differentiation biased populations during early and mid stages of recovery; Definition and application of the Spermatogonial Differentiation Index clarified. RNA sequencing analysis section updated to incorporate reanalysis of previously published single-cell RNA sequencing datasets, strengthening conclusions regarding stage-specific SPRY4-dependent transcriptional programs, including regulation of Id1 and Cxcl12; Discussion updated to integrate these findings with niche interaction and stem cell maintenance models. Section on MAPK/ERK signaling revised to clarify the role of ERK subcellular localization, distinguishing cytoplasmic versus nuclear ERK activity and its relationship to mTORC1 signaling during spermatogonial regeneration; Pharmacologic inhibition experiments refined to better distinguish ERK-dependent versus mTORC1-dependent effects. Long-term regeneration and fertility analyses expanded to include delayed recovery of undifferentiated spermatogonia, extended fertility assessment, and reduced sperm counts at later time points; New data added describing age-associated depletion of undifferentiated spermatogonia in Spry4 knockout males. Figures 1 through 5 revised for clarity and consistency with updated analyses; Multiple supplemental figures added or reorganized to support new data and improve presentation; Figure legends updated throughout; References updated and minor text edits made for clarity and consistency; Author affiliations and acknowledgments updated.

