## Supplemental Figures for "Restoration of Spermatogenesis is Dependent on Activation of a SPRY4-ERK Checkpoint Following Germline Stem Cell Damage"

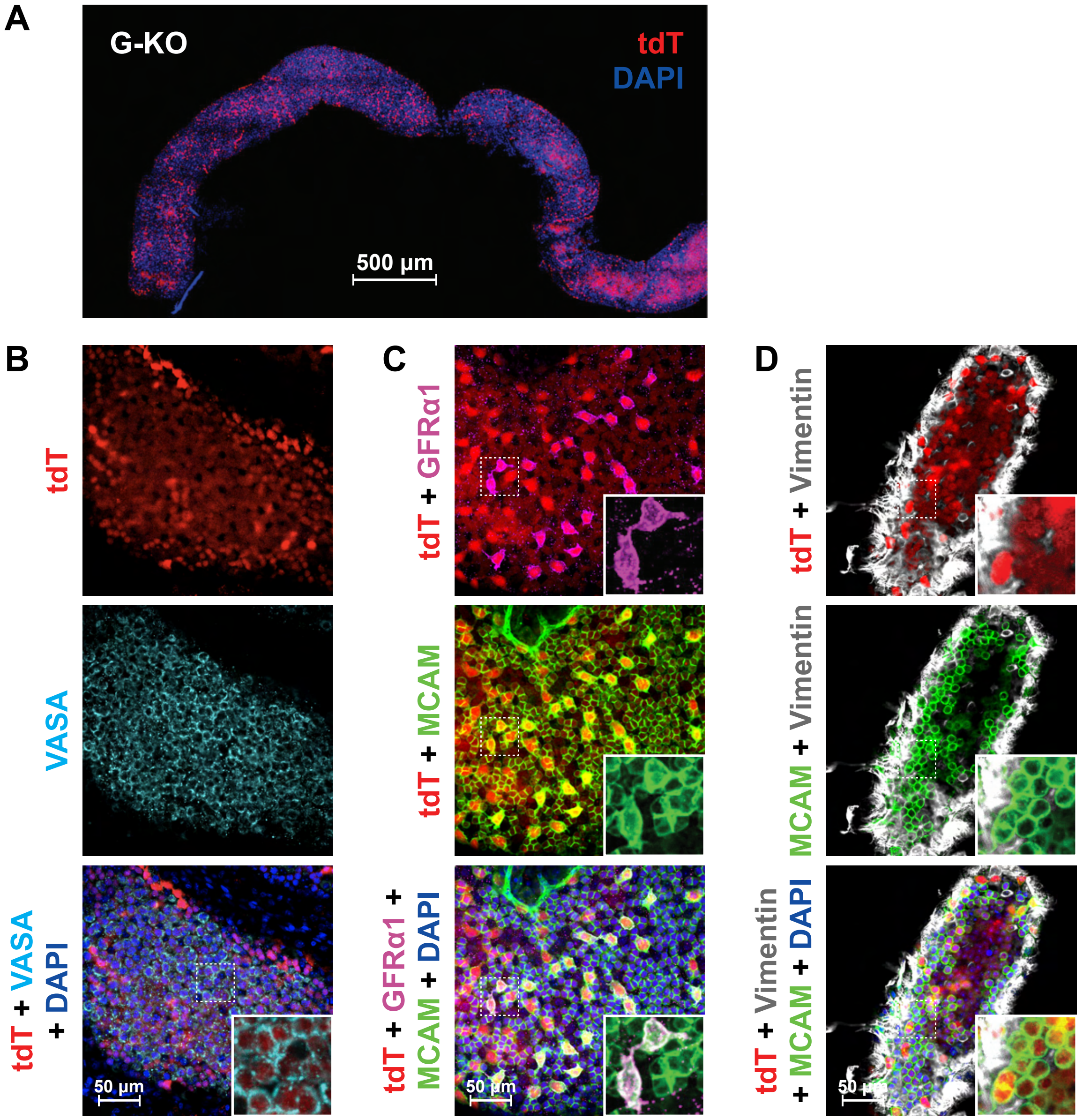


**Supplementary Figure 1. Fluorescent reporter in the *Spry4*$^{G\text{-}KO}$ testis labels germ cells of all stages.**

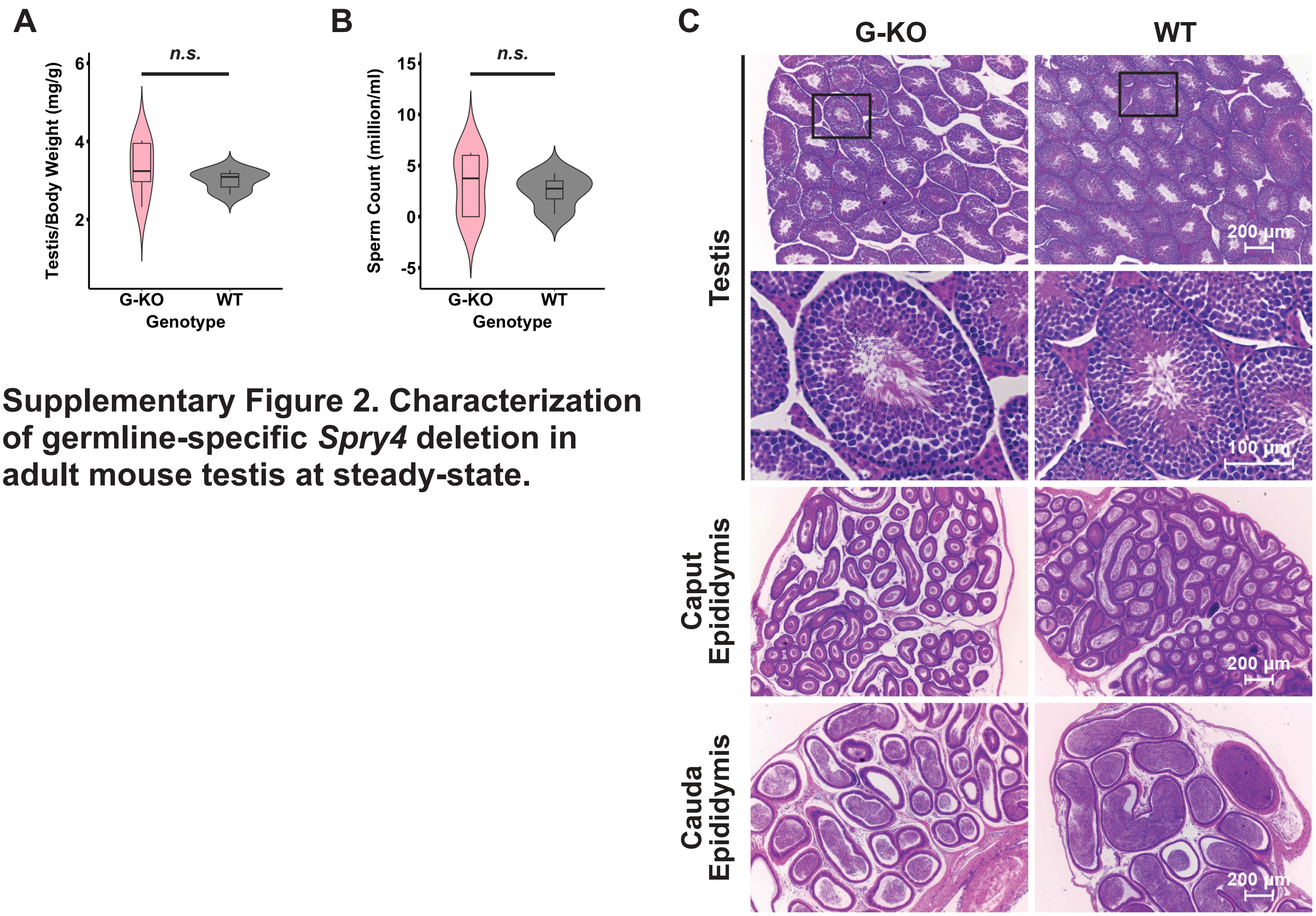


**Supplementary Figure 2. Characterization of germline-specific *Spry4* deletion in adult mouse testis at steady-state.**

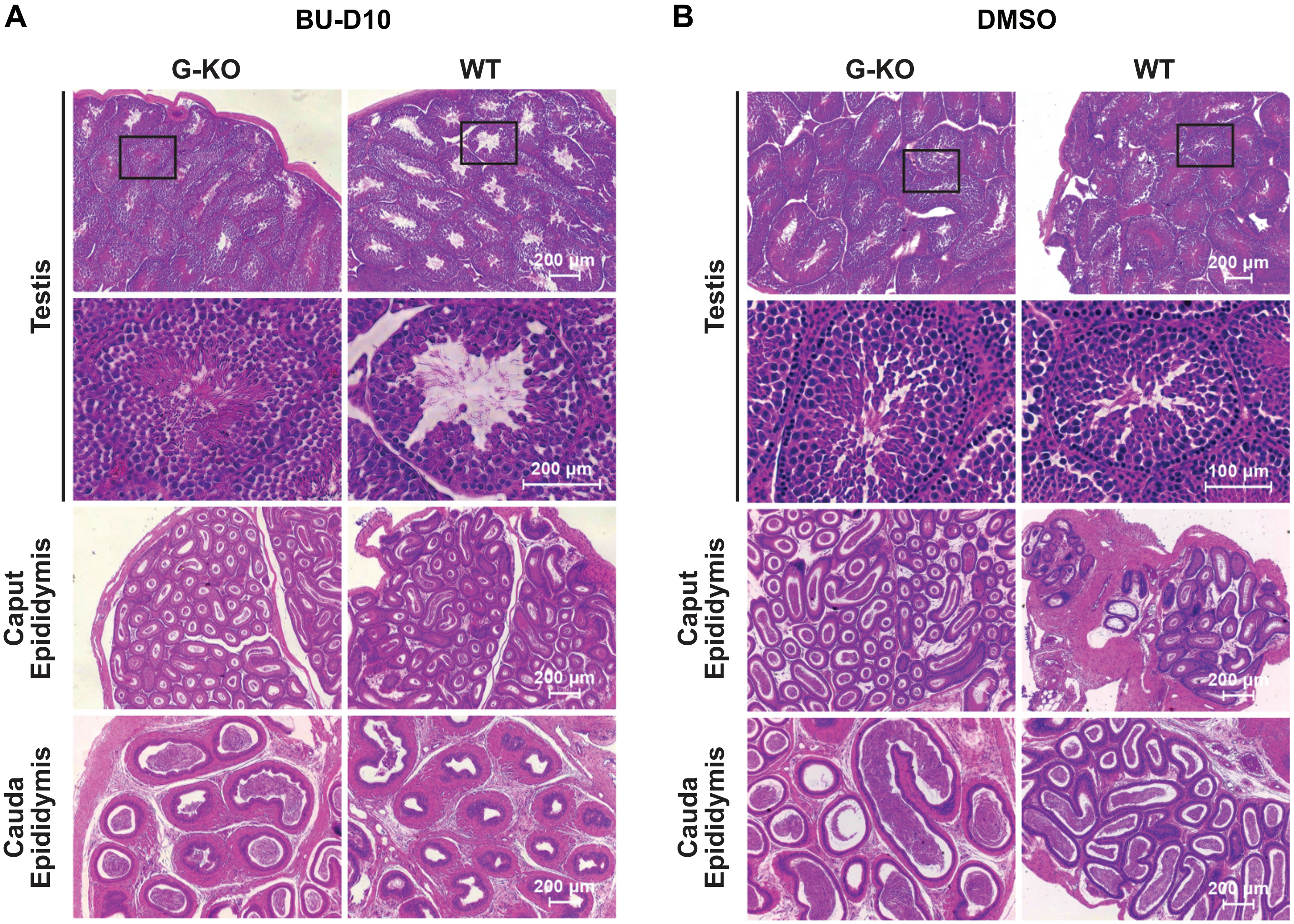


**Supplementary Figure 3. H&E-stained histological cross-sections of mouse testes and epididymides.**

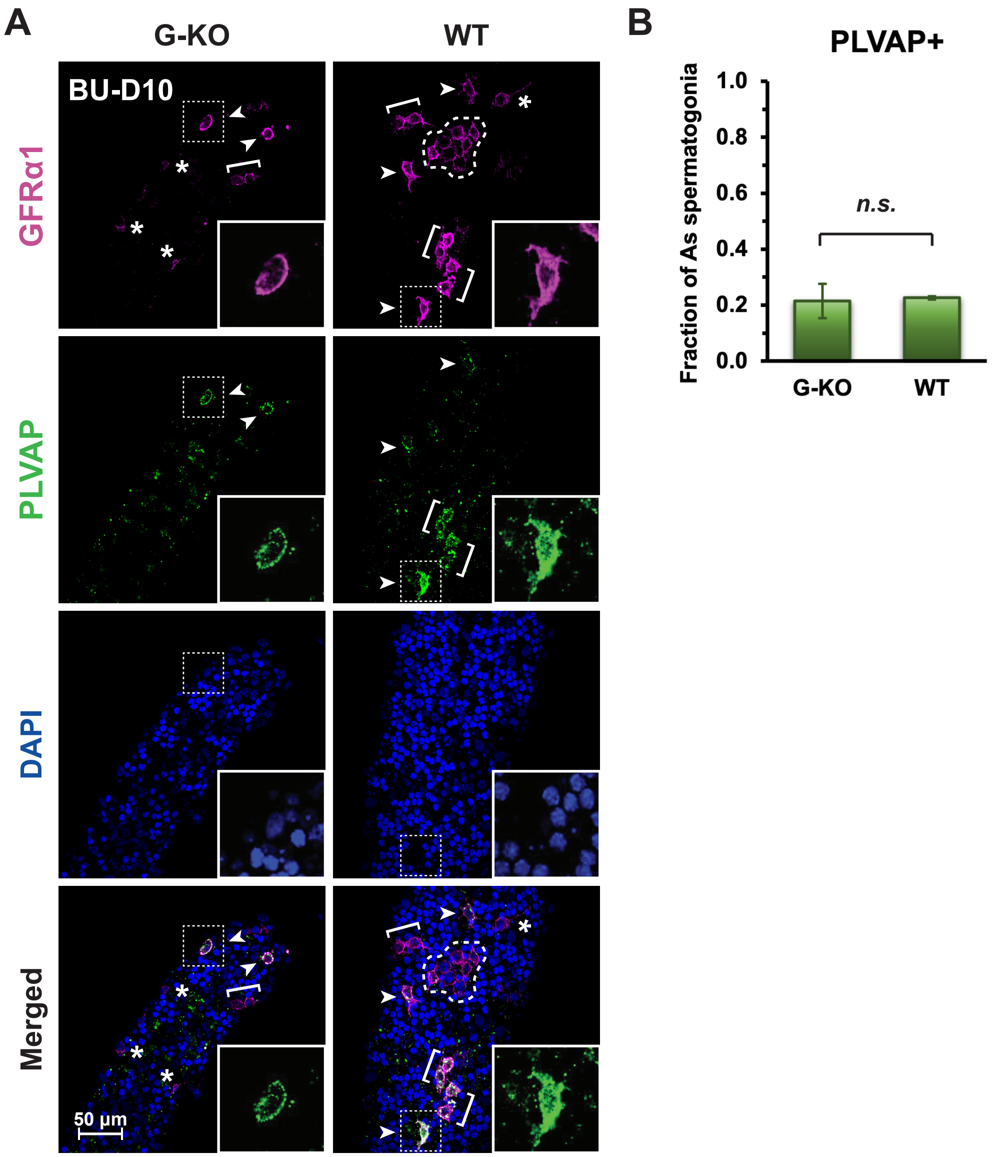


**Supplementary Figure 4. Characterization of $A_{undiff}$ spermatogonia with PLVAP.**

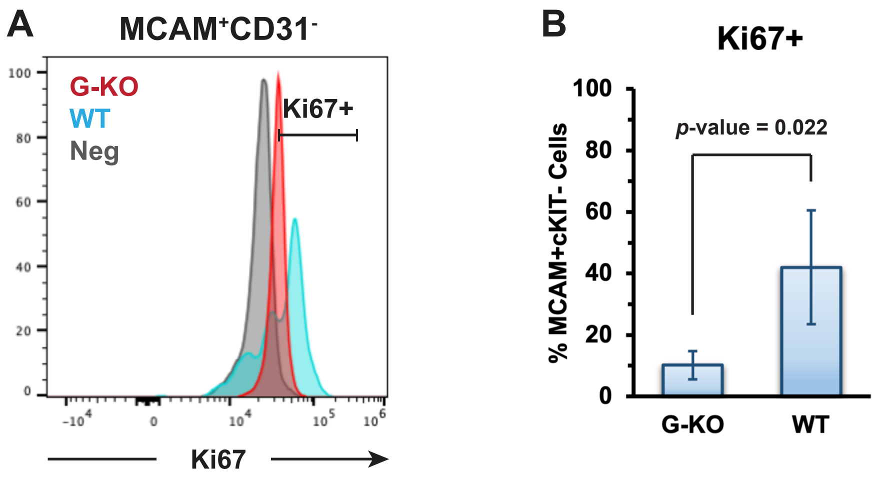


**Supplementary Figure 5. Flow cytometry analysis of MCAM$^{+}$ spermatogonia at BU-D10.**

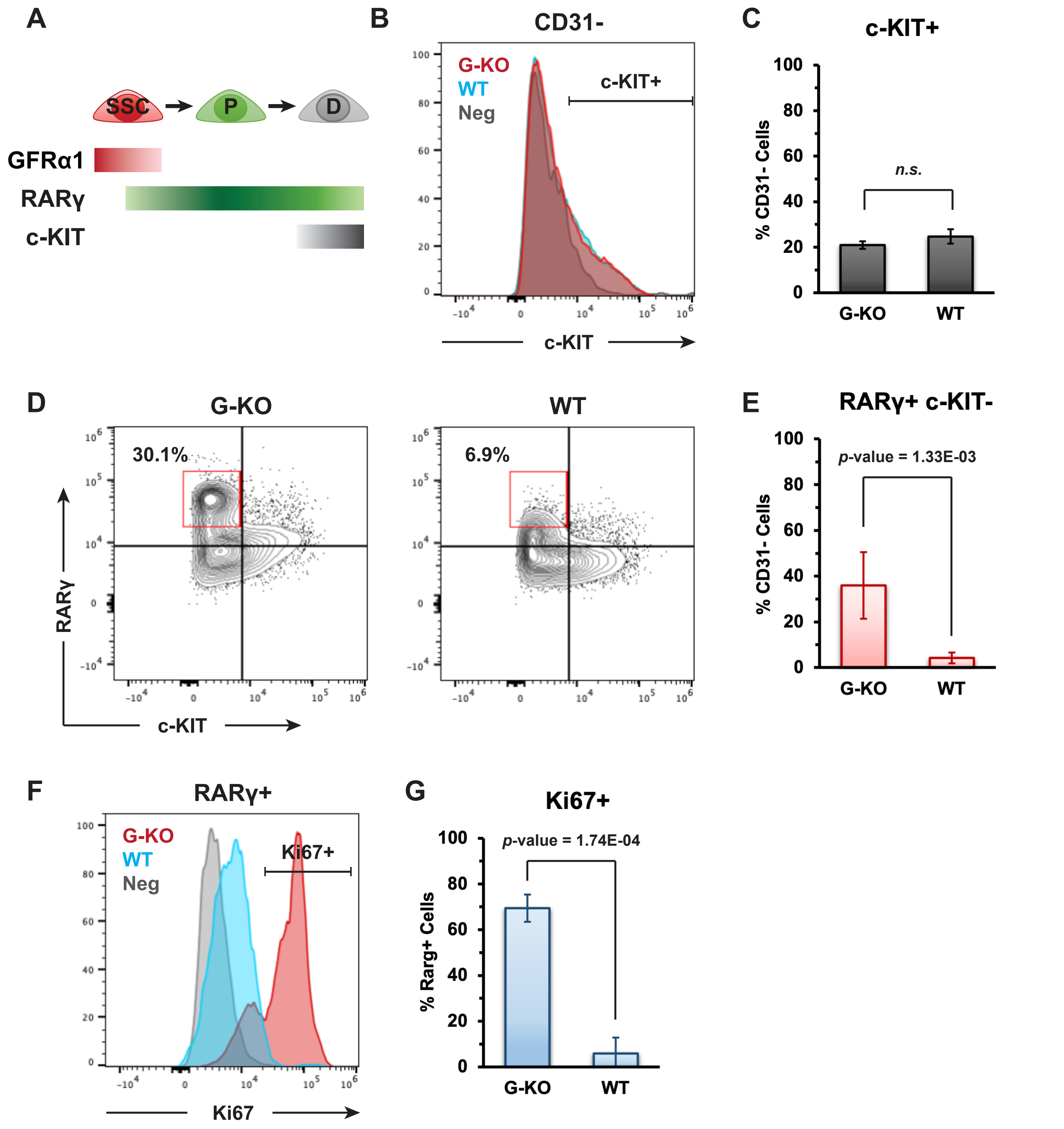


Supplementary Figure 6. Flow cytometry analysis of $A_{diff}$ spermatogonia at BU-D4.

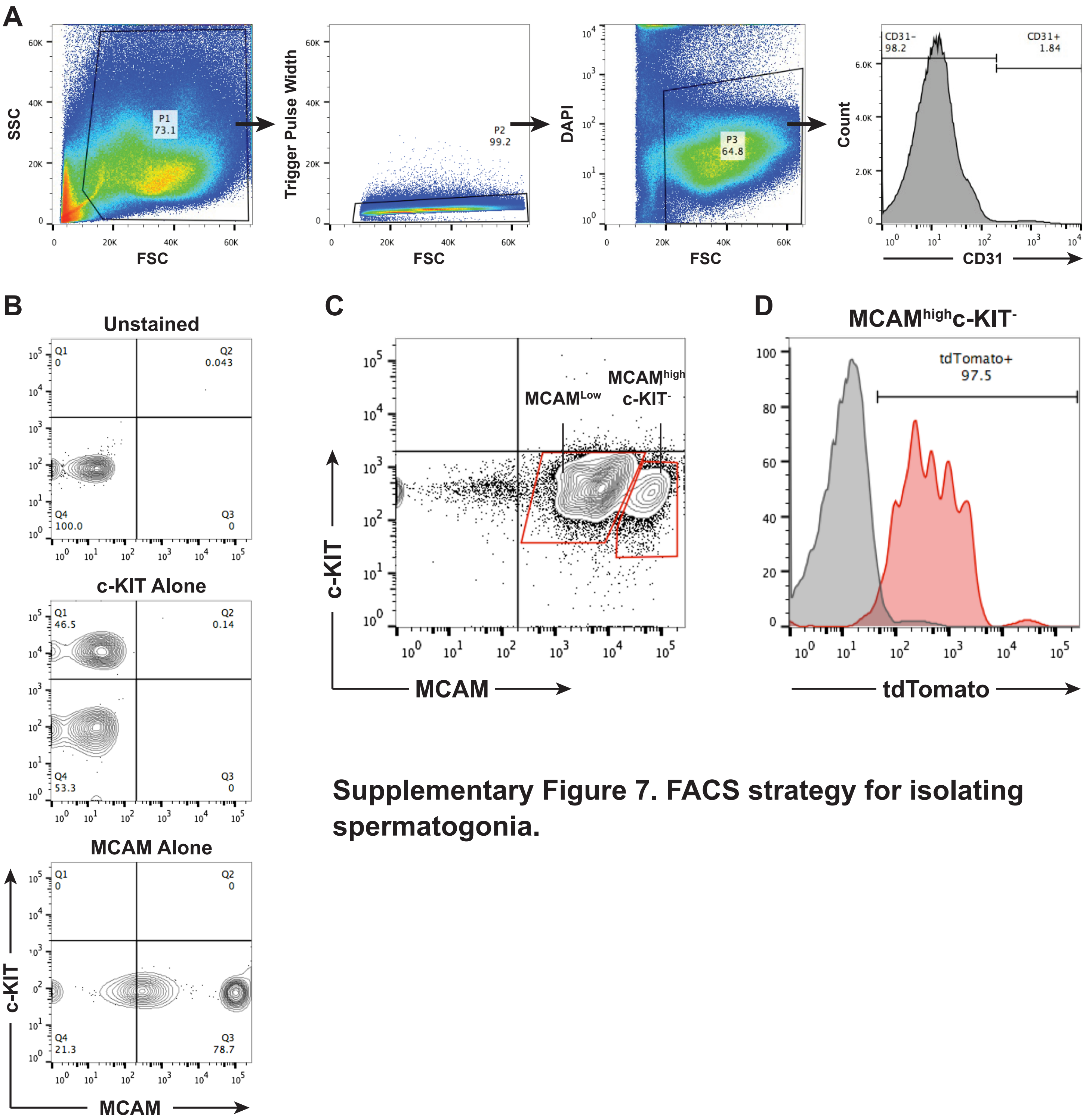


**Supplementary Figure 7. FACS strategy for isolating spermatogonia.**

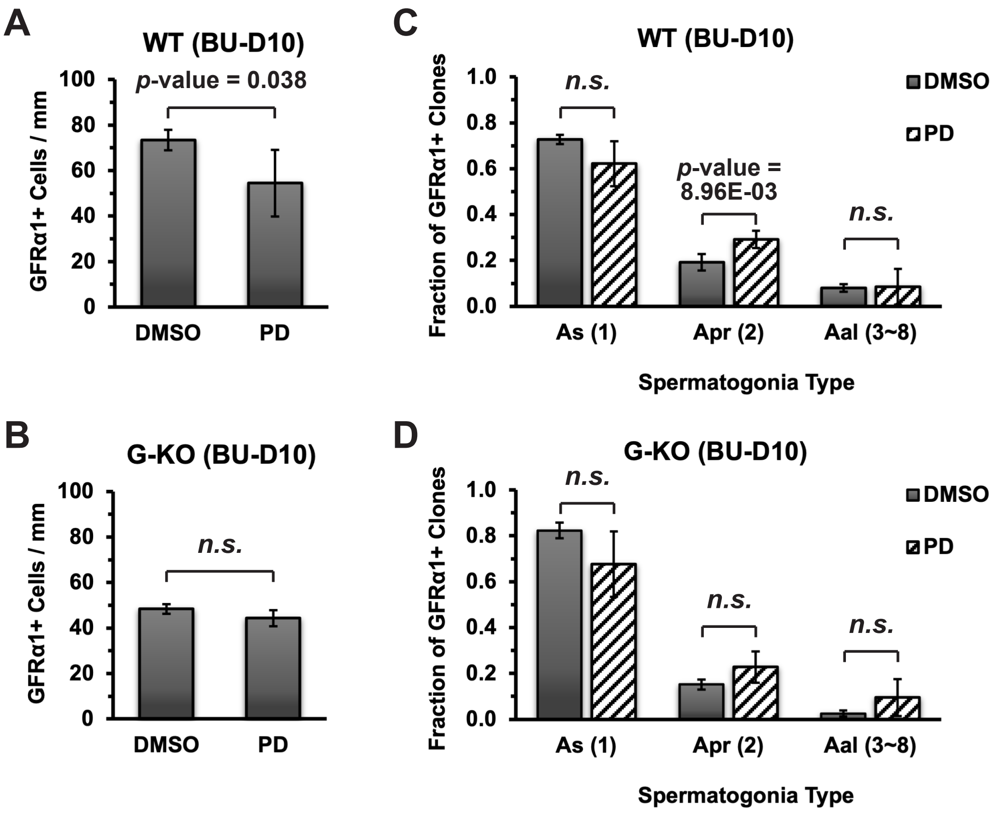


**Supplementary Figure 8. Characterization of $A_{undiff}$ spermatogonia with PD0325901 (PD) treatment.**

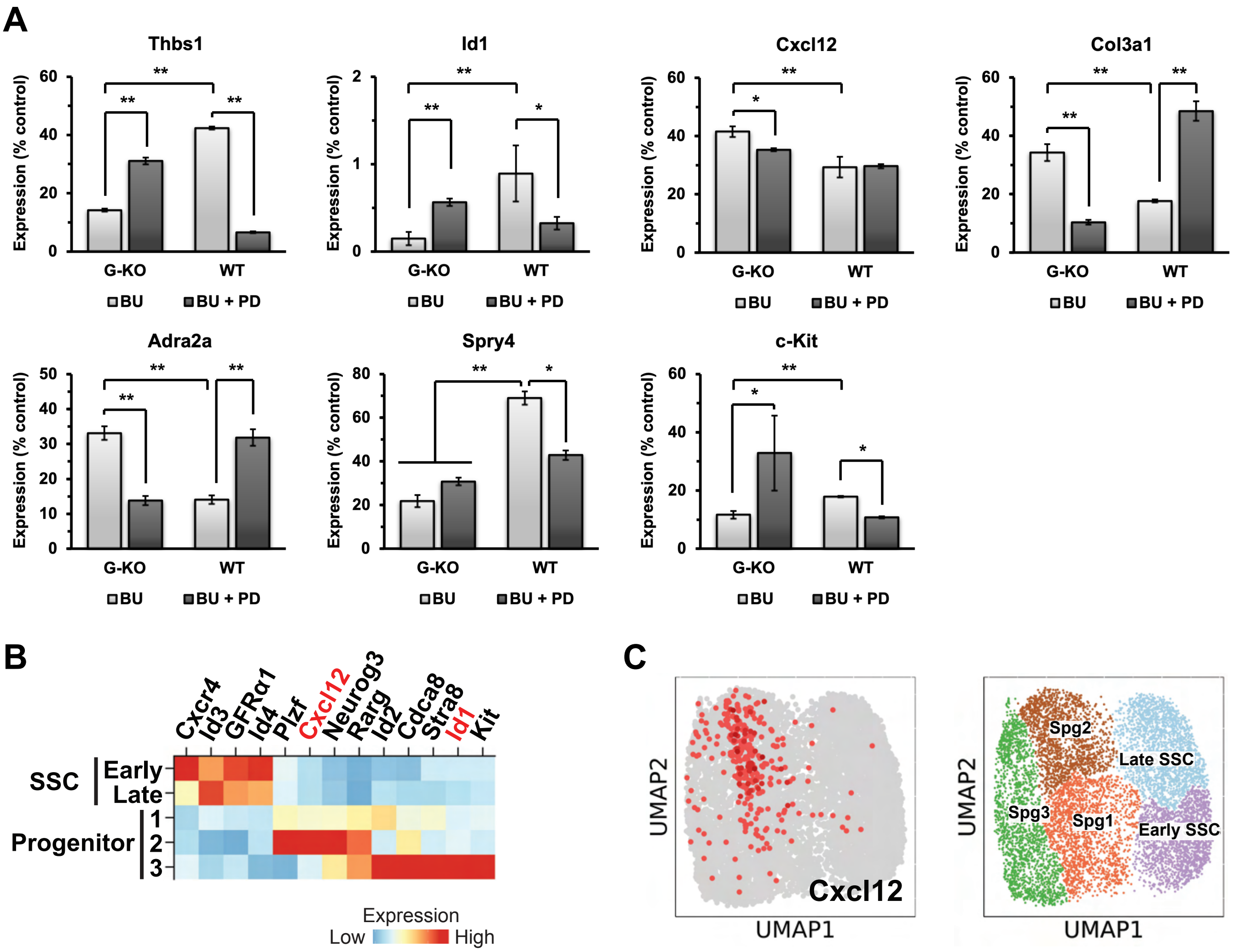


**Supplementary Figure 9. Expression of selected genes *in vivo*.**

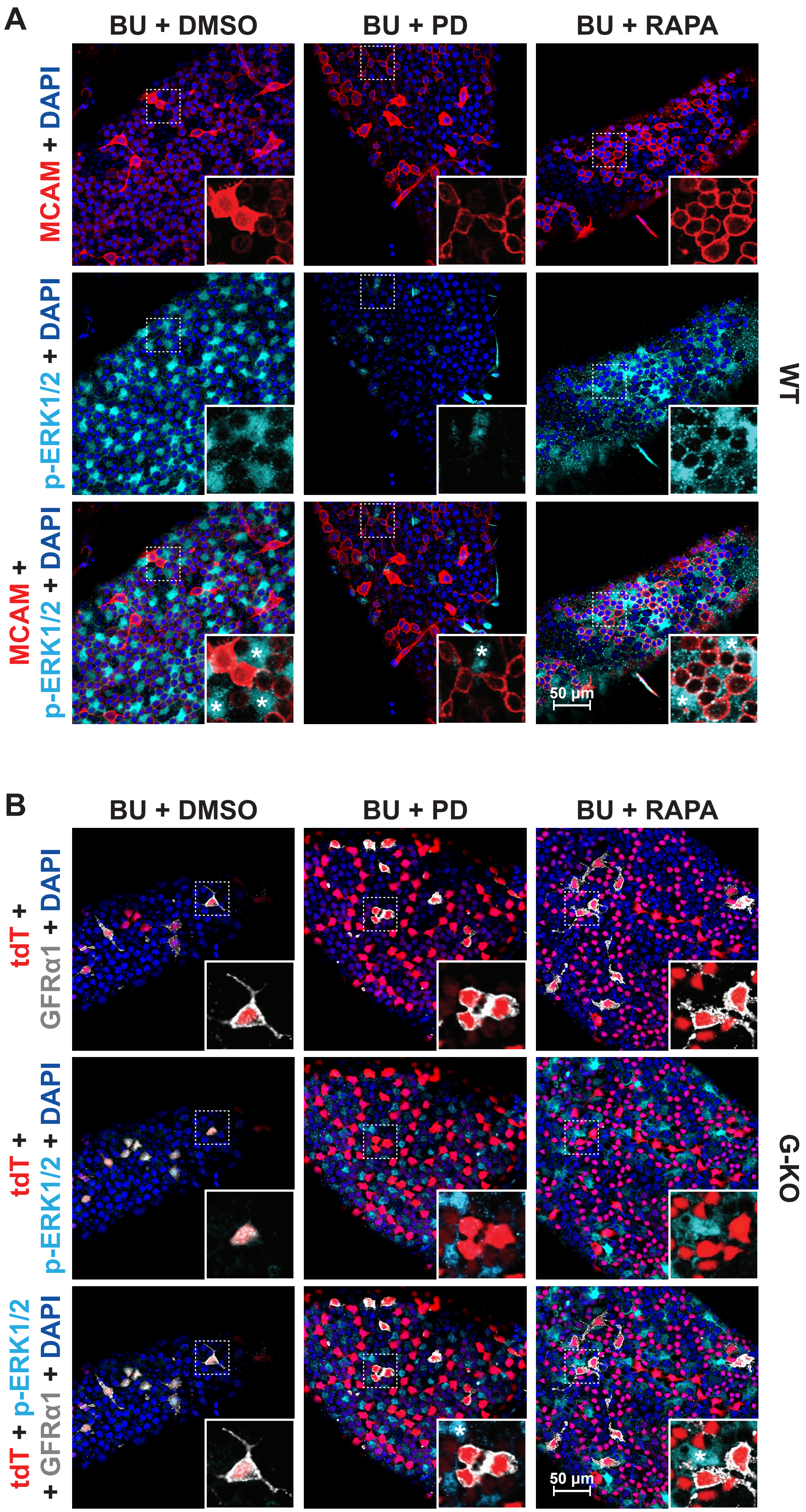


**Supplementary Figure 10. ERK activity within the seminiferous tubules during regeneration (BU-D10).**

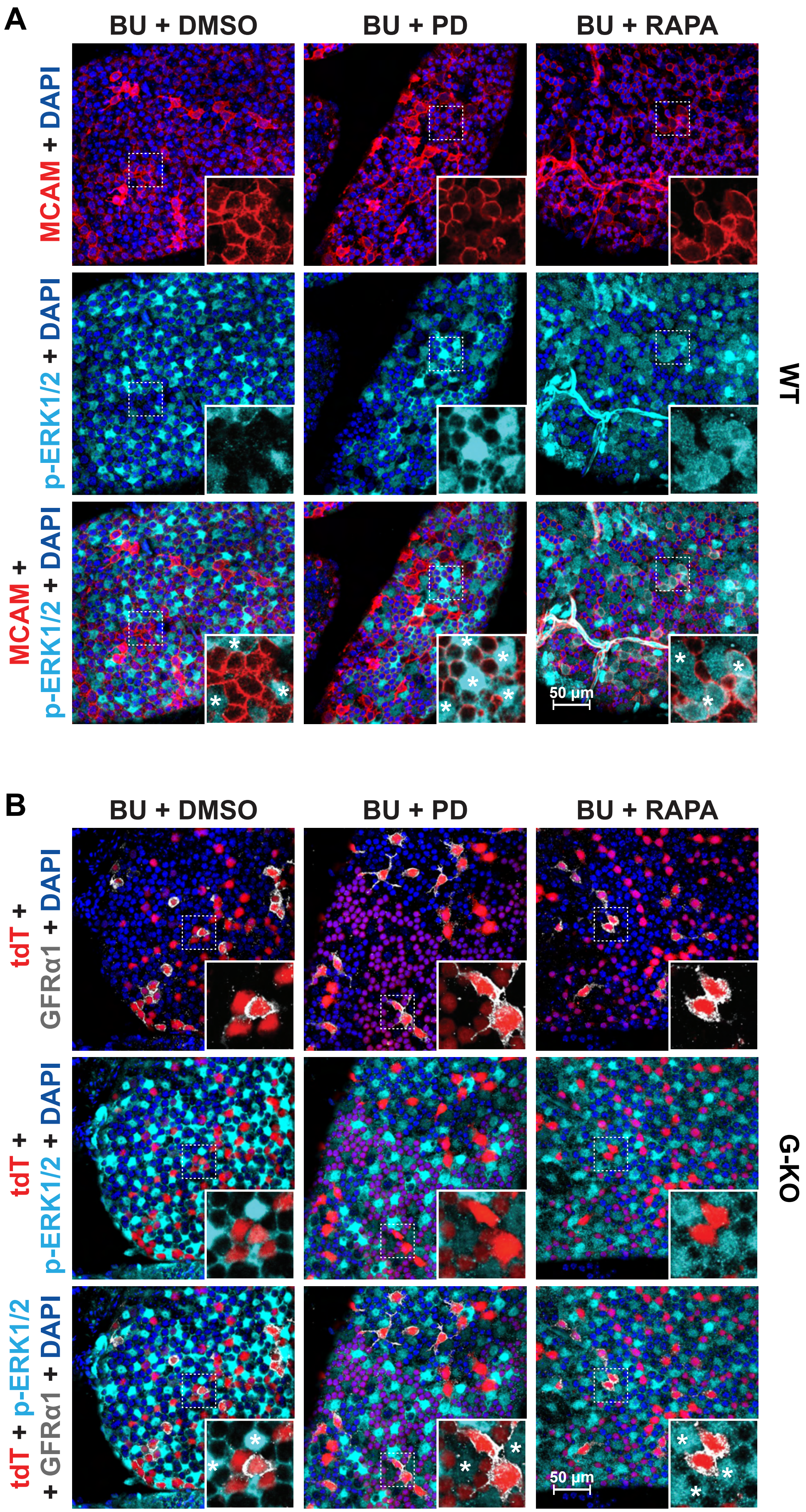


**Supplementary Figure 11. ERK activity within the seminiferous tubules quickly after damage (BU-D2).**

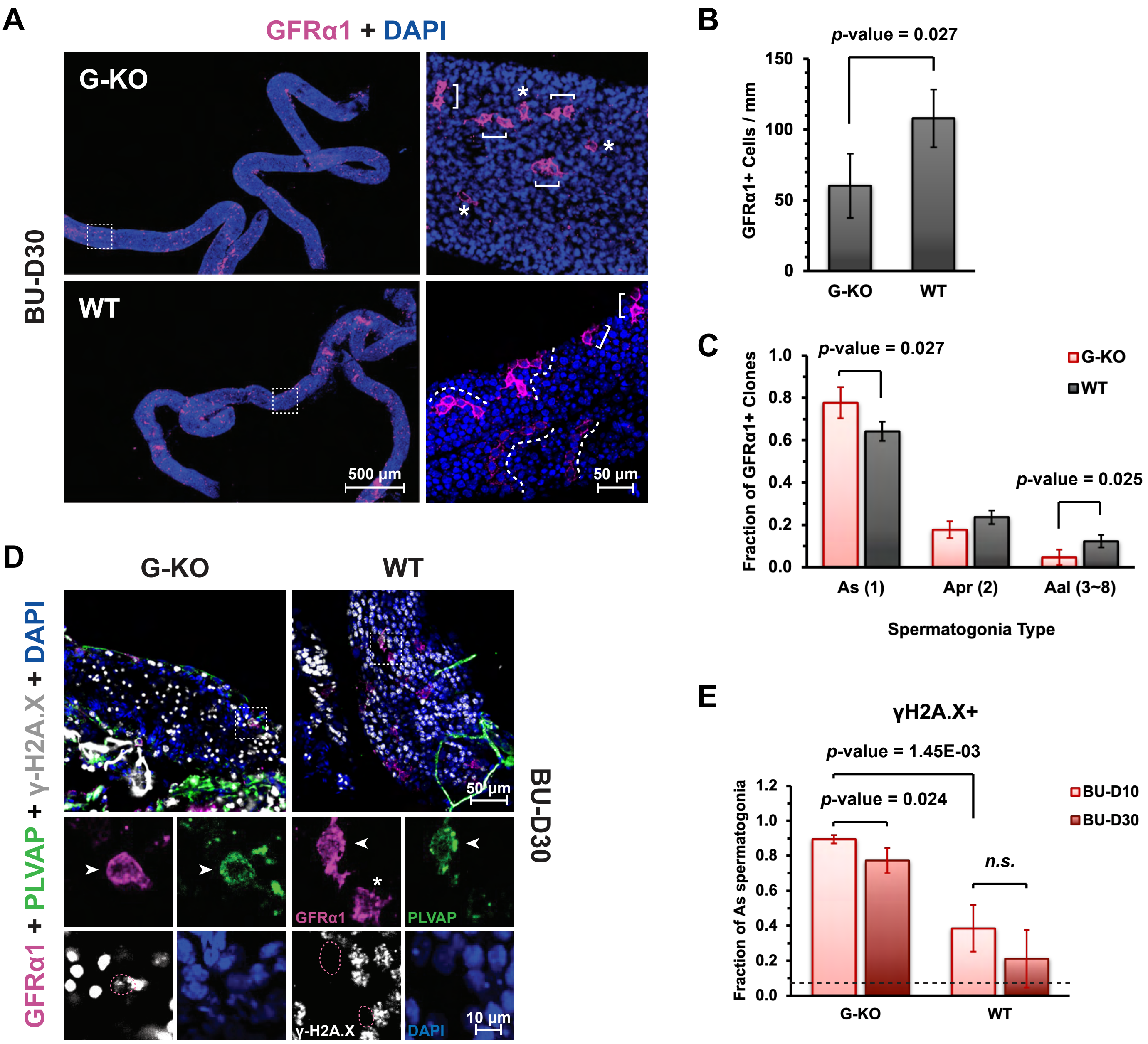


**Supplementary Figure 12. Spermatogonia remain reduced with intensive DNA damage in adult *Spry4*$^{G\text{-}KO}$ mice 30 days after damage (BU-D30).**

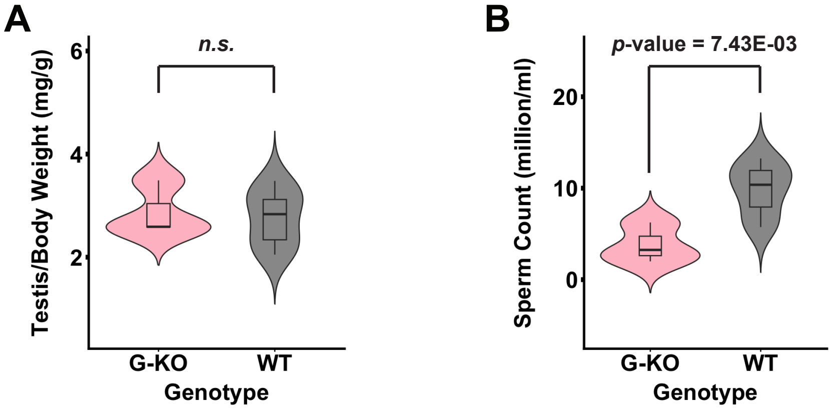


**Supplementary Figure 13. Characterization of adult mouse testes recovered long-term after damage.**

- H&E: G-KO vs. WT ?

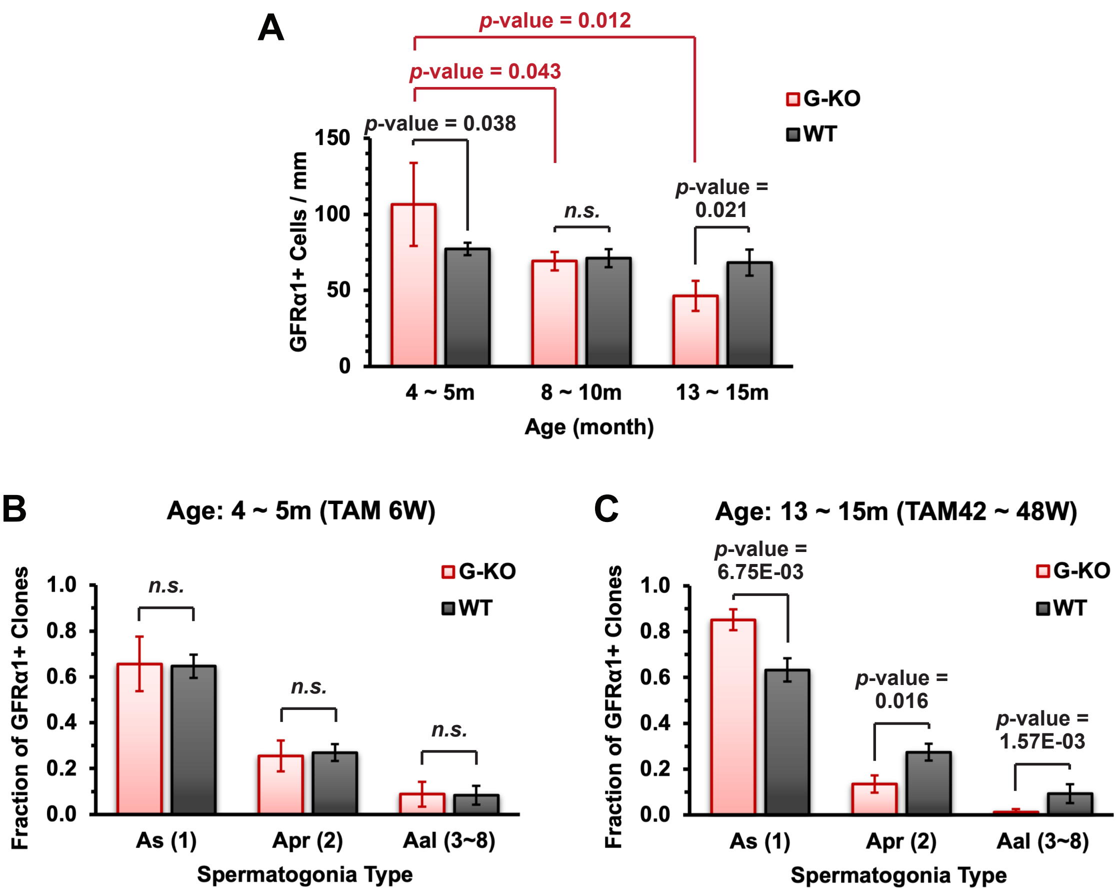


**Supplementary Figure 14. $A_{undiff}$ spermatogonia in adult *Spry4*$^{G\text{-}KO}$ mice are progressively reduced and enriched to the most primitive $A_s$ spermatogonia (SSCs) in aging males.**
